# Induction of Recurrent Break Cluster Genes in Neural Progenitor Cells Differentiated from Embryonic Stem Cells In Culture

**DOI:** 10.1101/2019.12.30.891168

**Authors:** Aseda Tena, Yuxiang Zhang, Nia Kyritsis, Anne Devorak, Jeffrey Zurita, Pei-Chi Wei, Frederick W. Alt

## Abstract

Mild replication stress enhances appearance of dozens of robust recurrent genomic break clusters, termed RDCs, in cultured primary mouse neural stem and progenitor cells (NSPCs). Robust RDCs occur within genes (“RDC-genes”) that are long and have roles in neural cell communications and/or have been implicated in neuropsychiatric diseases or cancer. We sought to develop an *in vitro* approach to determine whether specific RDC formation is associated with neural development. For this purpose, we adapted a system to induce neural progenitor cell (NPC) development from mouse embryonic stem cell (ESC) lines deficient for XRCC4 plus p53, a genotype that enhances DNA double-strand break (DSB) persistence to enhance detection. We tested for RDCs by our genome wide DSB identification approach that captures DSBs genome-wide via their ability to join to specific genomic Cas9/sgRNA-generated bait DSBs. In XRCC4/p53-deficient ES cells, we detected 7 RDCs, which were in genes, with two RDCs being robust. In contrast, in NPCs derived from these ES cell lines, we detected 29 RDCs, a large fraction of which were robust and associated with long, transcribed neural genes that were also robust RDC-genes in primary NSPCs. These studies suggest that many RDCs present in NSPCs are developmentally influenced to occur in this cell type and indicate that induced development of NPCs from ES cells provides an approach to rapidly elucidate mechanistic aspects of NPC RDC formation.

**SIGNIFICANCE STATEMENT:** We previously discovered a set of long neural genes susceptible to frequent DNA breaks in primary mouse brain progenitor cells. We termed these genes RDC-genes. RDC-gene breakage during brain development might alter neural gene function and contribute to neurological diseases and brain cancer. To provide an approach to characterize the unknown mechanism of neural RDC-gene breakage, we asked whether RDC-genes appear in neural progenitors differentiated from embryonic stem cells in culture. Indeed, robust RDC-genes appeared in neural progenitors differentiated in culture and many overlapped with robust RDC-genes in primary brain progenitors. These studies indicate that *in vitro* development of neural progenitors provides a model system for elucidating how RDC-genes are formed.

## INTRODUCTION

The DNA Ligase 4, and its obligate XRCC4 co-factor, are core C-NHEJ factors that each required for mouse lymphocyte development, due to essential roles in V(D)J recombination (1, 2). Moreover, they are each required for neural development (2, 3). Absence of either of these factors led to wide-spread p53-dependent death of newly generated neurons due to the inability to repair DSBs generated in neuro-progenitors (4, 5). While p53 deficiency rescued XRCC4-deficient neuronal apoptosis, the p53/XRCC4-double-deficient mice routinely died from medulloblastomas (MBs) with recurrent genomic rearrangements characteristic of those found in the sonic hedgehog sub-group of this human childhood brain cancer (6, 7). The striking overlaps of these findings with those obtained for the same mutant genotypes with respect to effects on development of lymphocytes and cancers of progenitor lymphocytes led to speculation that specific DSBs may impact neural development or disease (8, 9, 10). However, identification of such putative breaks was challenging with available technologies. To elucidate potential recurrent DSBs in developing neural progenitors, we employed the linear amplification-mediated, high-throughput, genome-wide, translocation sequencing (LAM-HTGTS) to map, at nucleotide resolution, DSBs in primary XRCC4/p53 deficient NSPCs based on ability to join to bait DSBs introduced by Cas9/sgRNAs (11). Our initial LAM-HTGTS experiments on *ex vivo* propagated mouse neural stem and progenitor cells (NSPCs) employed HTGTS bait DSBs on 3 different mouse chromosomes and identified 27 recurrent DSB clusters that, strikingly, were nearly all in bodies of long neural genes and mostly only observed after treatment with aphidicolin (APH) to induce mild replication stress (11).

To extend these studies, we exploited our finding that joining of two DSBs occurs more frequently if they lie on the same *cis* chromosome (12, 13). Thus, we introduced DSBs into each of the 20 mouse chromosomes as baits for HTGTS libraries from control or APH-treated NSPCs (14). This analysis confirmed previously identified RDCs and identified many more. Again, most RDCs were in genes and were identified upon NSPC treatment with APH to generate mild replication stress. Based on identification frequency with different baits and RDC-DSB density, we ranked relative RDC-gene robustness. On this basis, 19 originally identified plus 11 newly identified RDC-genes were highly robust (14). Of note, four of these highly robust RDC-genes were identifiable in untreated controls, but became more robust with APH-treatment, consistent with ectopic replication stress augmenting an ongoing endogenous process (11, 14). The great majority of highly robust RDC-genes are very long (>0.5Mb), variably transcribed, encode proteins that regulate synaptic function/cell adhesion, and have been associated with neuropsychiatric and development disorders and/or cancer (11, 14). We categorized such genes as Group 1 RDCs. We also identified Group 2 RDCs that contain multiple genes with at least one greater than 80kb long, and Group 3 RDCs which are clusters of small (<20kb) genes. Group 2 and 3 RDCs are mostly less robust than Group 1 RDCs and also less frequently associated with neuropsychiatric and neurodevelopmental disorders (14), and are not a focus of this study. Recently, LAM-HTGTS studies identified 36 DSB clusters in very long genes (analogous to group 1 RDCs) in human neural precursors cells derived from human-induced pluripotent stem cells, of which about 70% were orthologs of mouse RDC-genes (15).

Some RDCs overlap with common fragile sites (CFSs) and copy number variations (CNVs), that have been suggested to be fragile due to collisions between transcription and replication related processes in very long genes (11, 16, 17, 18). However, detailed mechanisms that cause RDC-gene DSBs in neural progenitor cells largely remain to be elucidated. For example, given the variable levels of overall robust RDC-gene transcription (11, 14), the precise role of transcription in RDC generation, or even if it is required, is an important, open question. Likewise, the question of when robust RDCs arise in the context of NSPC development and the factors, including stress, that promote their occurrence is also unknown. Addressing such questions experimentally in primary NSPCs that arise *in vivo* during mouse development is very challenging technically. Therefore, to facilitate mechanistic and developmental studies of RDC-gene formation in NSPCs, we sought to employ an *in vitro* system for induction of neural progenitor cell (NPC) differentiation from embryonic stem cells (ESCs) (19). Comparison of RDC-genes in ESCs to those in NPCs derived from them *in vitro* (ESC-NPCs) could provide insights into mechanisms that lead to robust RDC-occurrence during NPC development. Likewise, if RDCs arise *de novo* in such an *in vitro* differentiation system, targeted genetic modifications could be introduced to RDC-genes or their regulatory sequences in parental ESCs and assess effects on RDC formation subsequent to their induction to ESC-NPC or even to mature neurons in culture. Here, we report that robust RDC formation does indeed occur in ESC-NPCs generated form ESCs in culture.

## RESULTS

### HTGTS Bait DSBs to Identify RDCs in Embryonic Stem Cells

In our initial RDC experiments, we identified RDCs in primary NSPCs via introduction of Cas9/sgRNA mediated bait DSBs into specific sites on 3 mouse chromosomes, including chromosomes *12, 15* and *16* (11). To investigate if mouse ESCs also harbor RDCs, we used the same general strategy, with chromosomes 12 and 15 HTGTS baits as in the earlier experiments, along with a chromosome 7-specific bait DSB (*SI Appendix, Table S1*), instead of chromosome 16 HTGTS bait. For unknown reasons we did not achieve robust cutting with the chromosome 16-specific Cas9/sgRNA in ESCs. The Cas9/sgRNA constructs were individually introduced in *Xrcc4*^*−/−*^*p53*^*−/−*^ ESCs (referred to as ESC Line 1) on one chromosomal site at a time. Based on our previous studies, the *Xrcc4*^*−/−*^*p53*^*−/−*^ background leads to DSB persistence, which similarly to primary NSPC studies can facilitate detection of ESC RDCs (11, 14). In each experiment, ESCs were treated with APH to induce mild replication stress or with DMSO (vehicle control). Experiments with each bait DSB were repeated four times and analyzed as described (11). For HTGTS library data analyses, we applied our custom-designed RDC-pipeline (20), which identified 5 RDCs in the first tested ESC line (ESC line 1), and only appeared after APH treatment (Figs. 1*A-C*). These 5 RDCs were all located within genes (Figs. 1*D-H*). To corroborate these findings, we repeated the experiments in a second genotype-matched ESC line (referred to as ESC line 2) and identified 5 APH-induced RDCs (Figs. S1*A-C*) that partially overlapped (3 of 5) with the set in ESC line 1 (Figs. S1*D-H*).

**Figure 1:**
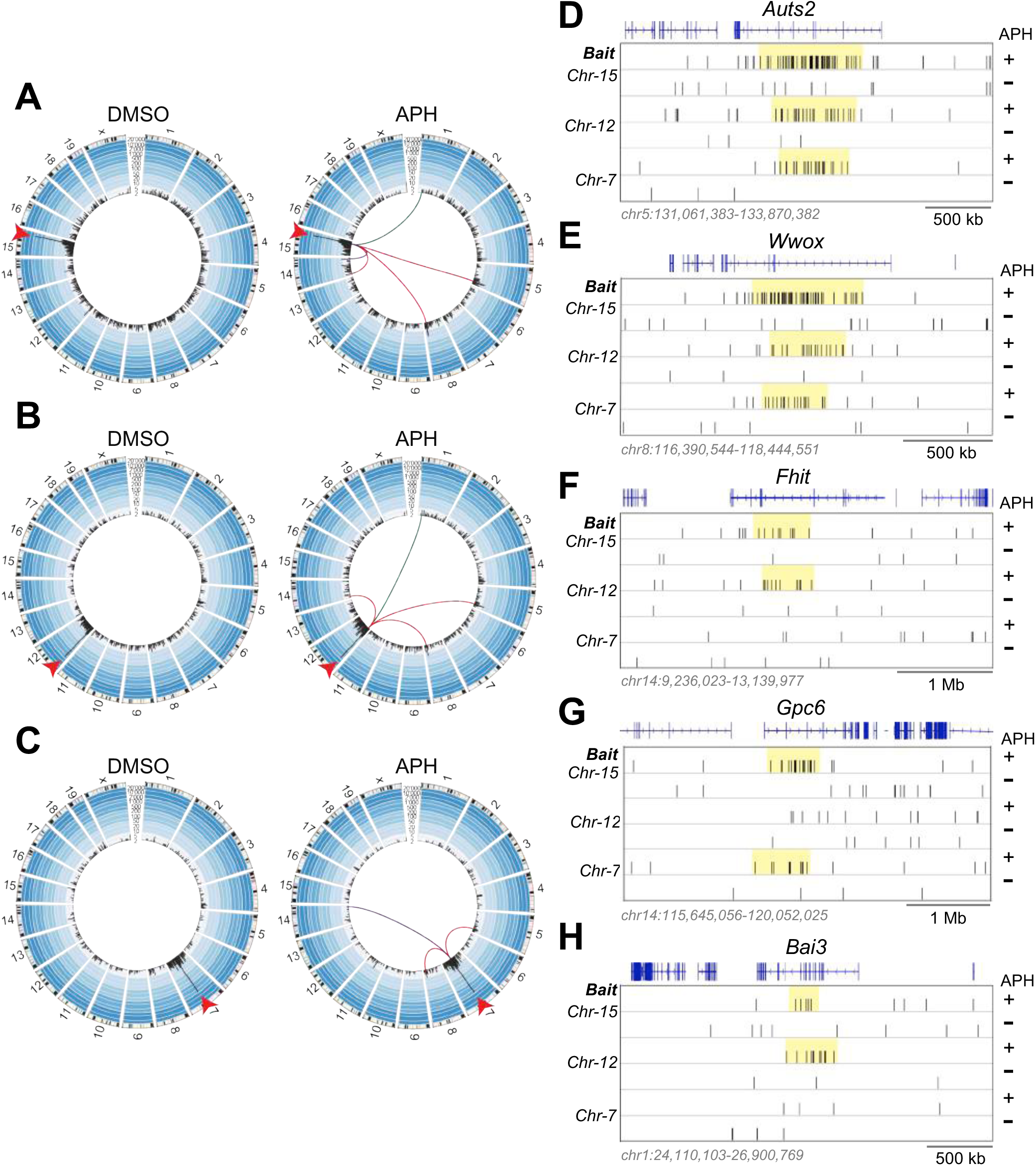
Genome-wide Identification of Replication Stress-induced RDC-genes in Embryonic Stem Cells. (***A-C***) Circos plots of the mouse genome divided into individual chromosomes show the genome-wide LAM-HTGTS junction pattern in *Xrcc4*^*−/−*^*p53*^*−/−*^ ESCs. Junctions identified by LAM-HTGTS baits locating at *Chr*-15, *Chr*-12, or *Chr*-7 were shown as black bars in per 2.5-Mb bin. Bar height indicates the number of translocations per bin on a log scale. 10,000 randomly selected junctions from four independent experiments are plotted in DMSO- (*Left* panels) and APH-treated cells (*Right* panels). Red arrowheads in circos plots denote the bait DSB site for each bait chromosome. Red lines in APH-treated experiments (*Right* panels) connect the break-site to 3 replication stress-induced RDCs identified by bait DSBs on all three tested chromosomes. The purple and green lines connect the break-site to RDCs identified by two of the three HTGTS baits. **(*D-H)*** 20,000 randomly selected LAM-HTGTS prey junctions from APH-(+) or DMSO-treated (-) ESCs are plotted. Panels represent 5 RDC-genes discovered by either two or three independent HTGTS-baits on the indicated chromosomes. The yellow rectangle indicates the RDC location. RefGene (blue track) indicates the gene location.

The 7 confirmed ESC RDCs all occurred within Group 1 RDC-genes (Figs.1*D-H*; Figs. S1*D-H*), with lengths ranging from 0.5Mb-1.6Mb (*SI Appendix*, Table S4). Three ESC RDC-genes (*Auts2, Wwox* and *Fhit*) were shared between both lines, but only *Auts2* and *Wwox* were classified as robust RDCs based on high RDC DSB junction density and the fact that they were captured by all three different HTGTS-baits, (Figs. 1*D,E*; Figs. S1*D,E*). *Fhit*, along with the four ESC RDCs that were unique to each line, were considered weaker RDCs (lower DSB junctions and discovered by only two of the three baits), (Figs. 1*F-H;* Figs. S1*F-H*). Notably, the two unique RDCs in the ESC line 1(*Gpc6* and *Bai3*) were both RDC-candidates in ESC line 2 (i.e. found with one HTGTS bait); while *Dock1* a unique RDC in ESC line 2 was a candidate in ESC line 1 (*SI Appendix*, Dataset S2). We note that the majority of RDCs identified in both ESC lines (6 of 7) were identified in *trans* by DSB baits employed on chromosomes 12, 15 and 7 (Figs. 1*D-H;* Fig. S1*D-F* and *H*). Only *Dock1*, an RDC-gene in ESC line 2 resided on a bait-chromosome (Fig. S1*G*). We also note that four RDCs *(Auts2, Gpc6, Wwox and Fhit*) overlapped with previously reported CNVs in ESCs (17), supporting the notion that RDC DSBs may contribute to formation of CNVs *in Xrcc4*^*−/−*^*p53*^*−/−*^ *ESC lines* (21). Furthermore, these RDCs have been mapped as common fragile sites in human fibroblast lines (17, 22, 23). Finally, all 7 RDCs identified in ESCs were identified as RDCs in prior NSPC studies (11, 14) and four (*Auts2, Wwox, Gpc6* and *Bai3*) were also RDC-genes in ESC-NPCs (*See below*). *Dock1*, was an RDC-candidate in both ESC-NPC lines (*SI Appendix*, Dataset S3).

### Identification of RDCs in embryonic stem cell–induced neural progenitor cells

We differentiated both *Xrcc4*^*−/−*^*p53*^*−/−*^ ESC line 1 and 2 into *Xrcc4*^*−/−*^*p53*^*−/−*^ NPCs to test whether this process would lead to generation of NPCs that had formed additional RDCs compared to those in their parental ESCs. For both *Xrcc4*^*−/−*^*p53*^*−/−*^ ESC lines, we employed immunofluorescence assays to confirm the differentiation of *Xrcc4*^*−/−*^*p53*^*−/−*^ ESCs into NPCs based on lack of expression of an ESC-specific *Oct4* and gain of expression of the NPC-specific *Sox1* and *Nestin* markers (Fig. S2*A*). We further tested *Xrcc4*^*−/−*^*p53*^*−/−*^ ESC-derived NPCs (*Xrcc4*^*−/−*^*p53*^*−/−*^ ESC-NPCs) by measuring their ability to give rise to neurons in a monolayer differentiation culture system (19). After 10 days in culture medium supplemented with retinoic acid, *Xrcc4*^*−/−*^*p53*^*−/−*^ ESC-NPCs downregulated *Sox1* and gained expression of neuronal-specific markers *NeuN* and *β-III tubulin* (Fig. S*2B*). To test the ability of *Xrcc4*^*−/−*^*p53*^*−/−*^ ESC-NPCs to generate RDCs, we performed HTGTS based on Cas9/sgRNA specific bait DSBs introduced into chromosomes *12, 15 and 7*, as described above for ESC HTGTS experiments. Each experiment also was done with or without APH treatment, repeated four times, and analyzed as previously described (11, 14). For *Xrcc4*^*−/−*^*p53*^*−/−*^ NPCs derived from ESC line 1, we identified 22 RDCs (*SI Appendix*, Dataset 1), which, as for primary NSPC RDCs, were enhanced by replication stress (Figs. 2*A-C*) and were located within genes (Figs. 2*D-G*). In addition, we performed a second set of studies with *Xrcc4*^*−/−*^*p53*^*−/−*^ NPCs derived from ESC line 2, which identified 24 RDCs (*SI Appendix*, Dataset 1), which were again enhanced by replication stress (Figs. S2*C-E*) and all located in genes (Figs. S2*F-I*).

**Figure 2:**
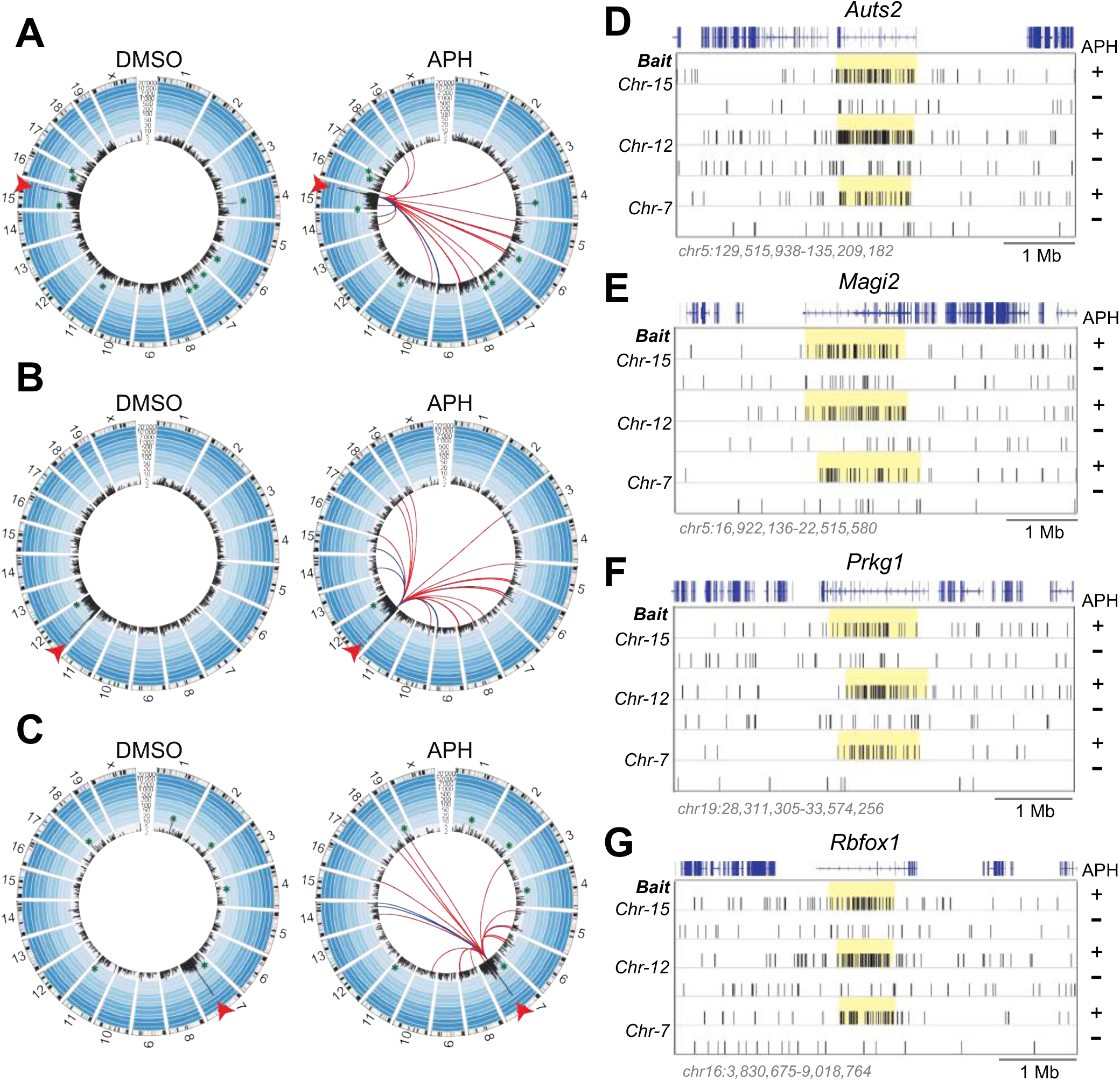
Genome-wide Identification of Replication Stress-induced RDC-genes in Embryonic Stem Cell-derived Neural Progenitor Cells. (***A-C***) Circos plots of the mouse genome divided into individual chromosomes show the genome-wide LAM-HTGTS junction pattern of *Chr-15, Chr-12*, and *Chr-7* sgRNA-mediated bait DSBs in *Xrcc4*^*−/−*^*p53*^*−/−*^ ESC-NPCs. Plots are organized as described in Figs. 1*A-C*. 10,000 randomly selected junctions from four independent experiments are plotted for DMSO- (*left* panels) and APH-treated (*right* panels) for each LAM-HTGTS bait experiment. Red lines in APH-treated experiments (*right* panels) connect the break-site to 17 replication stress-induced RDCs identified by bait DSBs on all three tested chromosomes. Blue lines connect break-site to RDCs identified by bait DSBs on two of the three tested break-sites, which numbered four for *Chr-15*, four for *Chr-12*, and two for *Chr-7* bait DSBs. LAM-HTGTS bait DSB sites (red arrowhead) and Cas9:sgRNA off-target sites (green asterisk) are denoted in both DMSO and APH plots. **(*D-G*)** 20,000 randomly selected LAM-HTGTS prey junctions from APH- (+) or DMSO-treated (-) experiments are plotted for *Auts2, Magi2, Prkg1* and *Rbfox1* RDC-genes. The yellow rectangle indicates the RDC location. RefGene (blue track) indicates the gene location.

Between both *Xrcc4*^*−/−*^*p53*^*−/−*^ ESC-NPC cell lines, we identified 29 RDC-genes, of which 17 appeared in *Xrcc4*^*−/−*^*p53*^*−/−*^ NPCs derived from both parental *Xrcc4*^*−/−*^*p53*^*−/−*^ ESC lines (Figs. 2D*-G*; Figs. S2*F-I*; Figs. S3*A,B*). In addition, 5 RDCs were found only in *Xrcc4*^*−/−*^*p53*^*−/−*^ NPCs derived from ESC line 1, and 7 RDCs were found only in *Xrcc4*^*−/−*^*p53*^*−/−*^ NPCs derived from ESC line 2, (*Figs. S3C,D*). Notably, 3 of 5 NPC line 1-specific RDCs were RDC-candidates in NPC line 2, and 5 out of 7 NPC line 2-specific RDCs were RDC-candidates in NPC line 1 (*SI Appendix*, Dataset S3). Among these RDC-genes, 27 were Group 1 RDCs which resided in very long genes and mostly occupied late replication domains (*SI Appendix*, Table S4). The other two RDCs were classified as Group 2 (*Tpgs2/Celf4*), and Group 3 (*Ackr2/1700048O20Rik*) RDCs (*SI Appendix*, Table S4). Similar to RDCs identified in primary NSPCs (11, 14), DSBs in ESC-NPCs RDC-genes map broadly across the length of the RDC-gene transcription unit, (Figs. 2*D-G*; Figs. S2*F-I*; *Figs. S3A-D*). Of the 17 shared RDCs, 14 were highly robust in both ESC-NPCs lines (*Fig. 3A; Fig. S4A; SI Appendix*, Dataset 1), and among the 30 most robust RDCs-genes detected in primary NSPCs by employing baits from all chromosomes (14). These findings demonstrate that *Xrcc4*^*−/−*^*p53*^*−/−*^ NPCs differentiated from *Xrcc4*^*−/−*^*p53*^*−/−*^ ESCs in culture gain the ability to generate a robust subset of RDCs observed in *Xrcc4*^*−/−*^*p53*^*−/−*^ primary NSPCs. We also identified many RDC-candidates in both ESCs and ESC-NPCs based on junctions with one bait, with the majority of these weak RDC-candidates residing, as expected, on a bait-containing chromosome (*SI Appendix*, Dataset S2; S3). It is possible that additional ESC and ESC-NPC RDCs would be confirmed if we employ baits from all chromosomes for analyses; but we also note, in this regard, that such an analysis only increased the number of detected robust NSPC RDCs to 30 versus the 19 detected in a three HTGTS bait analysis (11, 14).

**Figure 3:**
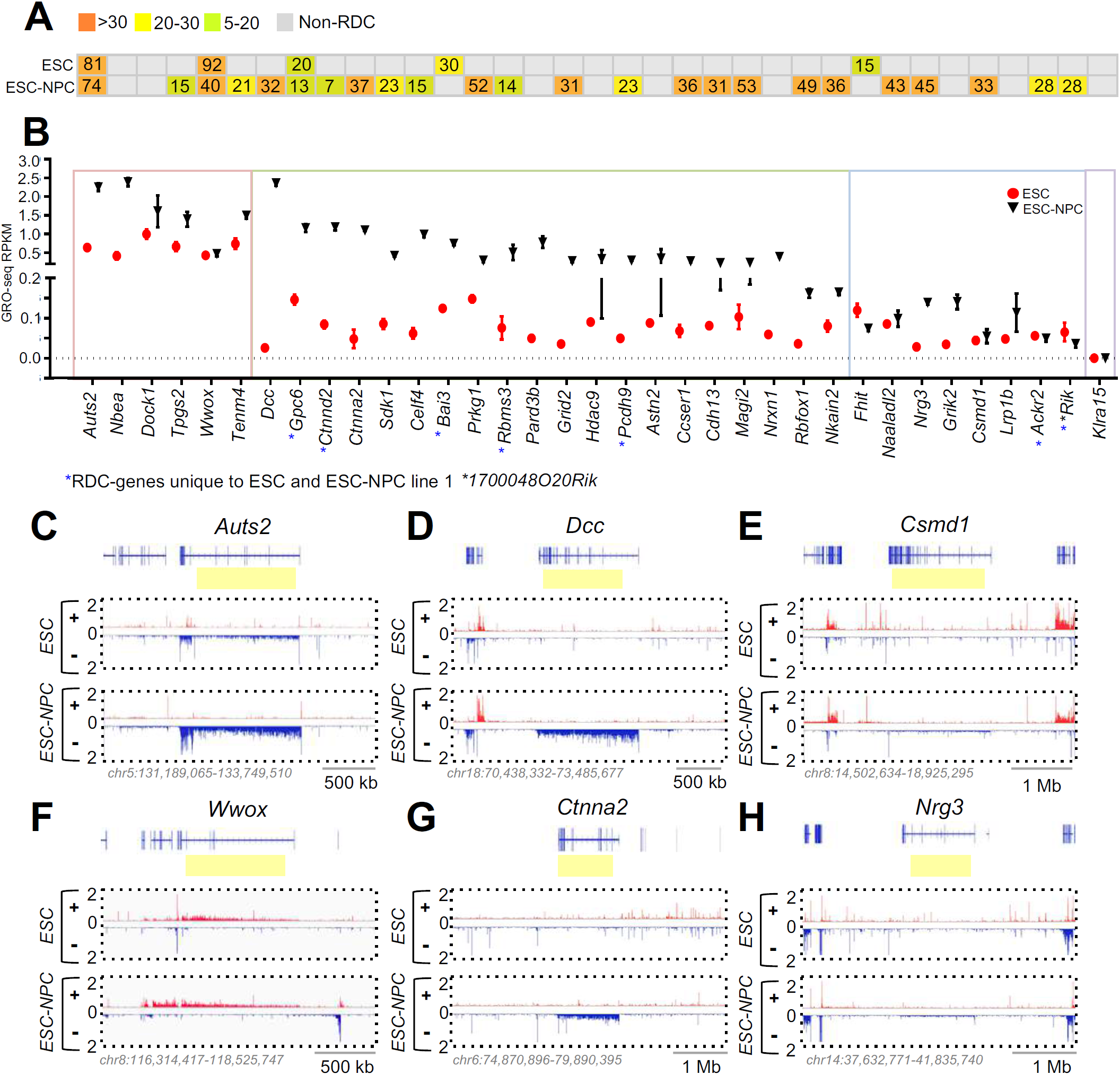
Transcription activity in embryonic stem cells and neural progenitor cells. ***(A)*** RDC robustness score (RRS) is indicated for each RDC-gene. Differences in RRS values (highest-lowest) are color-coated as illustrated. (***B***) Transcription activity measured by reads-per-kilobase-per-million (RPKM) from GRO-seq experiments is plotted for RDC-genes in both ESCs and differentiated NPCs. The most left box in the graph contains RDC-genes that have higher transcription activity in both ESCs and ESC-NPCs. The middle box contains RDC-genes that have higher transcription activity only in ESC-NPCs. The second right box contains genes with lower transcription activity in both cell types, and the last box contains *Klra15* which has no transcription activity in both cell types. Data represents (mean ± SD) based on 3 repeats. Blue asterisk: RDC-genes that are unique to cell line 1. **(*C-H*)** Panels represent robust RDC-genes in ESCs and differentiated NPCs with different transcription levels in both cell lines. Ordinate indicates normalized GRO-seq counts. Blue color indicates transcription orientation in centromeric-to-telomeric direction and red color telomeric-to-centromeric direction. RefGene (blue track) indicates the gene location and the yellow rectangle highlights the RDC location.

### Transcription activity of RDC-genes in embryonic stem cells and neural progenitor cells

Transcription patterns of RDC-genes found in *Xrcc4*^*−/−*^*p53*^*−/−*^ ESC lines versus RDC-genes found in *Xrcc4*^*−/−*^*p53*^*−/−*^ ESC-NPCs derived from them might provide clues to mechanisms underlying RDC-gene formation. We analyzed the transcription of all RDC-genes identified in *Xrcc4*^*−/−*^*p53*^*−/−*^ ESCs and *Xrcc4*^*−/−*^*p53*^*−/−*^ ESC-NPCs experiments via global run-on sequencing (GRO-seq). The 7 RDC-genes identified in *Xrcc4*^*−/−*^*p53*^*−/−*^ ESCs and the 29 RDCs found in *Xrcc4*^*−/−*^*p53*^*−/−*^ ESC-NPCs showed no consistent relationship between transcription levels and RDC formation. Some of the RDC-genes had high transcriptional activity in both cell types, others had high transcription activity only in *Xrcc4*^*−/−*^*p53*^*−/−*^ ESC-NPCs, while others had low transcription in *Xrcc4*^*−/−*^*p53*^*−/−*^ESCs and *Xrcc4*^*−/−*^*p53*^*−/−*^ ESC-NPCs (Fig. 3*B;* Fig. S4*B*). Moreover, the 14 robust *Xrcc4*^*−/−*^*p53*^*−/−*^ ESC-NPC RDCs also didn’t have a strong correlation between transcription levels and RDC-gene robustness score (Figs. 3*A,B;* Figs. S4*A,B*). More specifically, *Auts2* and *Wwox* are robust RDC-genes in both *Xrcc4*^*−/−*^*p53*^*−/−*^ ESCs and *Xrcc4*^*−/−*^*p53*^*−/−*^ ESC-NPCs, and have high transcription activity in both cell types (Fig. 3*B*, left box; Figs. 3*C,F*); *Dcc* and *Ctnna2* are *Xrcc4*^*−/−*^*p53*^*−/−*^ ESC-NPC specific RDCs with high transcription activity only in *Xrcc4*^*−/−*^*p53*^*−/−*^ ESC-NPCs and low transcription activity in *Xrcc4*^*−/−*^*p53*^*−/−*^ ESCs (Fig. *3B* middle box; Figs. 3*D,G*); while *Csmd1* and *Nrg3* are robust RDCs in ESC-NPCs despite their low transcription activity in these cells (Fig. 3*B* third quadrant; Figs. 3*E,H*). Similar examples are shown for the *Xrcc4*^*−/−*^*p53*^*−/−*^ ESC Line 2 and its derivative *Xrcc4*^*−/−*^*p53*^*−/−*^ ESC-NPCs (Figs. S4*A,B*; S*4C-H*). Thus, these ESC and ESC-NPC studies, together with our primary NSPCs studies (11, 14) indicate that transcription levels of RDC-genes *per se* have no obvious correlation with RDC formation in them.

## DISCUSSION

There are many fundamental questions related to molecular mechanisms that give rise to DSBs within RDC-genes in primary NSPCs that remain unanswered, in part due to difficulty in studying this phenomenon in primary cells from mice. We now show that induced development of NPCs from ESCs *in vitro* leads to the formation of a robust set of NPC specific RDC-genes, many of which overlap with robust RDC-genes in primary NSPCs. Therefore, the *in vitro* NPC differentiation system will offer a rapid approach to target and test genetic modifications in ESCs and then test their effects on robust RDC formation in ESC-NPCs. In some cases, mutations of interest could further be tested by differentiating the ESC-NPC cultures to neurons (19) or by using the ESCs for neural blastocyst complementation *in vivo* studies (24). The ESC-NPC differentiation studies showed that a large number of robust RDCs are absent in ESCs and only appear upon differentiation of ESCs into NPCs. Moreover, 26 of 36 replication-stress susceptible DSB ‘ hot spot’ genes identified in human-induced pluripotent stem cell-derived neural precursor cells were orthologs of primary mouse NSPC RDC-genes (11,14); and, 12 were orthologs of robust ESC-NPC RDCs identified in this study (15). These studies suggest that robust NPC RDCs are mostly cell type-specific and are developmentally influenced to occur in neural-specific genes in both human and mouse NPCs (15, this study). In this regard, further studies with ESC-NPC differentiation system to elucidate mechanisms underlying RDC formation should also be relevant to RDC formation in human NPCs.

We know that the majority of highly robust NSPC-RDCs and, now, ESC-NPC RDC-genes tend to arise in very long, transcribed neural genes associated with specific brain functions (11, 14, this study). Early studies performed on mouse ESCs and human fibroblasts found both spontaneous and replication-induced CNVs which were linked to long genes that were actively transcribed and late replicating (17, 21). A majority of mouse hotspot CNVs were also (CFSs) in human fibroblasts (22, 23). These studies led to speculation that mechanisms of breakage and CNV formation may involve collisions between the replication and transcription machinery that would be accentuated due to the long replication time of long genes (16, 17, 18, 25). Furthermore, because a subset of the robust primary NSPC and ESC-NPCs RDC-genes overlapped with mouse ESC CNVs and human fibroblast CFSs, it was proposed that this mechanism could potentially apply to RDC-gene formation (11, 14, 17). Yet, both our prior NSPC and current ESC-NPC RDC studies indicate that there is no direct relationship between robust RDC occurrence and the relative level of RDC-gene transcription. However, it is possible that very low transcription is sufficient for robust RDC formation. We can now directly test this notion by disrupting transcription of both highly and weakly transcribed robust RDC-genes in the *in vitro* NPC differentiation system and, if warranted by results, employ additional gene targeted mutational approaches to further test the potential roles of transcription. Studies could also be readily envisioned to employ the NPC *in vitro* differentiation approach to test other factors related to robust ESC-NPC RDC formation including potential roles of gene length, replication timing, and replication stress.

## MATERIALS AND METHODS

### Cell culture and LAM-HTGTS Bait DSB Induction

We used two genotype-matched, *de novo* derived embryonic stem cells that were *Xrcc4*^−/−^*p53*^−/−^ for the described studies. ESC-NPCs were generated as described (19) with minor modifications. LAM-HTGTS bait DSB induction at *Chr-7, Chr-12* and *Chr-15* was performed as described in ref. 11, 14 and SI. Details of cell cultures and LAM-HTGTS bait-induction are provided in *SI Appendix, Materials and Methods.*

### LAM-HTGTS

LAM-HTGTS Libraries were prepared as described previously (11, 14, 20) and sequenced on Illumina Mi-Seq and Next-Seq. Reads from demultiplexed FASTQ files were aligned to the genome build mm9/NCBI37 through Bowtie2 and processed through the HTGTS pipeline (20). In each library, only unique junctions were preserved for the RDC identification. *SI Appendix*, Table S3 lists the number of junctions for each experiment.

### RDC Identification

A SICER-based, unbiased, genome-wide method and a MACS-based method were both applied to identify APH-induced RDCs in *Xrcc4*^*−/−*^*p53*^*−/−*^ ESCs and *Xrcc4*^*−/−*^*p53*^*−/−*^ ESC-NPCs as described previously (11). RDC robustness score (RRS) was calculated as previously described in (14). Details are also provided in *SI Appendix, Materials and Methods.*

### GRO-seq

GRO-seq libraries were prepared as previously described (26). Three experimental replicates were performed for both *Xrcc4*^*−/−*^*p53*^*−/−*^ ESC lines and *Xrcc4*^*−/−*^*p53*^*−/−*^ ESC-NPC lines. Libraries were sequenced on Illumina Hi-Seq and Next-Seq. Details provided in *SI Appendix, Materials and Methods.*

## ACKNOWLEDGEMENTS

This work was supported by Children’s Hospital Department of Medicine and Charles H. Hood Foundation grant. P-C W. was supported by Charles A. King Trust Postdoctoral Research Fellowship Program, Bank of America, N.A., Co-Trustees. Y Z. is supported by a Damon Runyon Fellowship Program. FWA is an Investigator of the Howard Hughes Medical Institute.

## SUPPORTING INFORMATION

### MATERIALS AND METHODS

#### Embryonic stem cell culture, neural progenitor cell differentiation and HTGTS Bait DSB Induction

In brief, ESCs (*Xrcc4*^*−/−*^*p53*^*−/−*^ genotype) were established from blastocysts that were isolated from *Trp53*^*tm1Brd*^*/Trp53*^*+*^::*Xrcc4tm*^*2.1Fwa*^*/Xrcc4*^*tm2Fwa*^::*Tg(Nes-cre)1Kln/0* female mice bred with the *Trp53*^*tm1Brd*^*/Trp53*^*tm1Brd*^::*Xrcc4*^*tm2.1Fwa*^*/Xrcc4*^*tm2Fwa*^::*Tg(Nes-cre)1Kln/0* male mice. The ESCs were cultured on irradiated mouse feeder fibroblasts (MEFs) in the DMEM medium (Corning, 10-013-CV) supplemented with 15% ESC-grade fetal bovine serum (Atlanta biological, S11550H), 20mM HEPES (Gibco, 15630), non-essential amino acids (Gibco, 11140), 100 U/ml Pen/Strep (Gibco, 15140-122), 0.1mM beta-mercaptoethanol and 1000 U/mL ESGRO recombinant mouse leukemia inhibitory factor (Millipore, ESG1107). 2×10^6^ ESCs were mixed with 2 ug of single Cas9:sgRNA expression vector pX330 (1) for nucleofection (Lonza 4D Nucleofector, P3 solution, mouse ESC program). Cells were treated with 0.2 μM APH (Sigma, A4487) or diluted DMSO (1:590 in culture medium) for 72 hours. APH concentrations were reduced to 0.1 μM for another 24 hours, and genomic DNA was extracted for HTGTS library preparation.

#### Neural progenitor cells differentiation protocol

For monolayer NPC differentiation, 1×10^5^ ESCs were plated onto 10ug/ml of poly-L-ornithine- (PLO; Sigma, P4957) and laminin- (LMN; Sigma, L2020-1MG) coated 6-well tissue culture plates in N2B27 medium. N2B27 is a 1:1 mixture of DMEM/F12 (Gibco, 11330057) and Neurobasal (Gibco, 21103049) medium supplemented with 1% B27 minus vitamin A (Gibco, 12587-010), 0.5% modified N2 (25 μg/ml insulin, 100 μg/ml apo-transferrin, 6 ng/ml progesterone, 16 μg/ml putrescine, 30 nM sodium selenite and 50 μg/ml bovine serum albumin fraction V) and 0.5 mM Glutamax (Gibco, 35050061). After 6 days, cells were dissociated by accutase (StemCell Tech, 07920) and plated onto fresh PLO/LMN-coated 60mm plates in neural progenitor maintenance medium (NBBG) consisting of Neural Basal A (Gibco, 10888-022), 2% B27 (without retinyl acetate), 0.5mM Glutamax, 10μg/ml Gentamycin (Gibco, 15710-064),10ng/ml mouse FGFb (PHG0314, Invitrogen), 10ng/ml mouse PDGFbb (PMG0044, Invitrogen), and 10ng/ml human EGF (PHG0314, Invitrogen). The complete NPC induction was achieved after 6 days culture in NGGB. For each nucleofection experiment, 5×10^6^ NPCs were mixed with 5μg of pX330 Cas9:sgRNA plasmid and mouse neural stem cell reagent (Lonza, VPG-1004). NPCs were nucleofected using A033 program in the Nucleofector IIb (Lonza). NPCs were treated with 0.5 μM APH (Sigma) or diluted DMSO (1:590) DMSO for 72 hours. APH concentration was reduced to 0.25 μM for another 24 hours, and genomic DNA was extracted for HTGTS library preparation.

#### Neuron differentiation

For *in vitro* NPC-neuron differentiation, 4×10^5^ NPCs were seeded onto PLO/LMN-coated 12 mm coverslips. Cells were maintained in the neural differentiation medium (N2B27 supplemented with 25mM retinoic acid). Medium was refreshed daily. Cells were fixed at day 10 for immunocytochemistry staining.

#### Immunofluorescence staining

ESCs, ESC-NPCs and ESC-NPC-derived neurons were fixed in 4% paraformaldehyde at room temperature for 10 minutes. Cells were washed with DPBS three times and permeabilized by 0.5% Triton-X-100 for 5 minutes at room temperature. Cells were then incubated in the blocking solution (5% v/v goat serum, 0.5% w/v fish gelatin, 0.2% w/v Tween-20) for one hour, and then incubated with primary antibody probing for Pax6 (1:200, BioLegend, poly19013), Oct4 (1:400, Novus, GT486), Nanog (1:400, Abcam, ab80892), Nestin (1:400, Abcam, GR3175728-1), Sox1 (1:400, Cell Signaling, 41945), NeuN (1:400, Millipore, MAB377), and β-Tubulin III (1:1000, Abcam, ab18207). Cells were washed in 0.2% (v/v) Tween-20/DPBS three times, then incubated with secondary anti-mouse or anti-rabbit antibody (1:500; Alexa Fluor 488 or 594, Thermo Fisher Scientific). We used Fluoromount-G (SouthernBiotech) to stain the nuclei and mount the stained coverslips. Slides were analyzed with a Zeiss confocal spinning disk microscope (Yokogawa CSU-X1, 3i Intelligent Imaging Innovations). Images were processed with SlideBook6 software (3i Intelligent Imaging Innovations).

#### GRO-seq

GRO-seq libraries were prepared as previously described (4). 5-8×10^6^ nuclei were used for per GRO-seq library. GRO-seq data were aligned to mouse genome build mm9/NCBI37 by Bowtie2 and non-redundant, uniquely mapped sequence reads were retained. *De novo* transcripts were identified and gene expression levels were estimated as previously described (4). We randomly selected 10 million GRO-seq sequences from ESCs or ESC-NPCs libraries for comparative illustrations shown in Fig. 3 and in supporting Fig. 4. We calculated RPKM based on the length of longest transcript information extracted from RefGene. The average RPKM value (mean ± SD) from three independent GRO-seq experiments are shown in Fig. 3, supporting Fig. 4 and supporting Table 4.

#### RDC Robustness Score

As described previously (3), we used multiple comparison testing corrected *P*-value and the number of *trans*-chromosomal HTGTS baits that successfully identified a given RDC to determine the RDC robustness score (RRS).

#### Replication Timing Analysis

A custom Python script (2) was used to calculate median replication timing ratios of genomic regions based on mouse ESC-NPC Repli-chip data (5, 6). We apply log value 0.5 as cutoff for early replicating genes and minus 0.5 for late replicating genes, respectively.

#### Identification of sgRNA Off-Target Sites

Translocations between Cas9:sgRNA on- and off-target DSBs were identified by the MACS2 algorithm as previously described (2). The off-target sequences must have ≥ 30% similarity to the on-target sgRNA sequences and be found at least twice in the replicate experiments.

## SUPPLEMENTARY FIGURE LEGENDS

**Figure S1.**
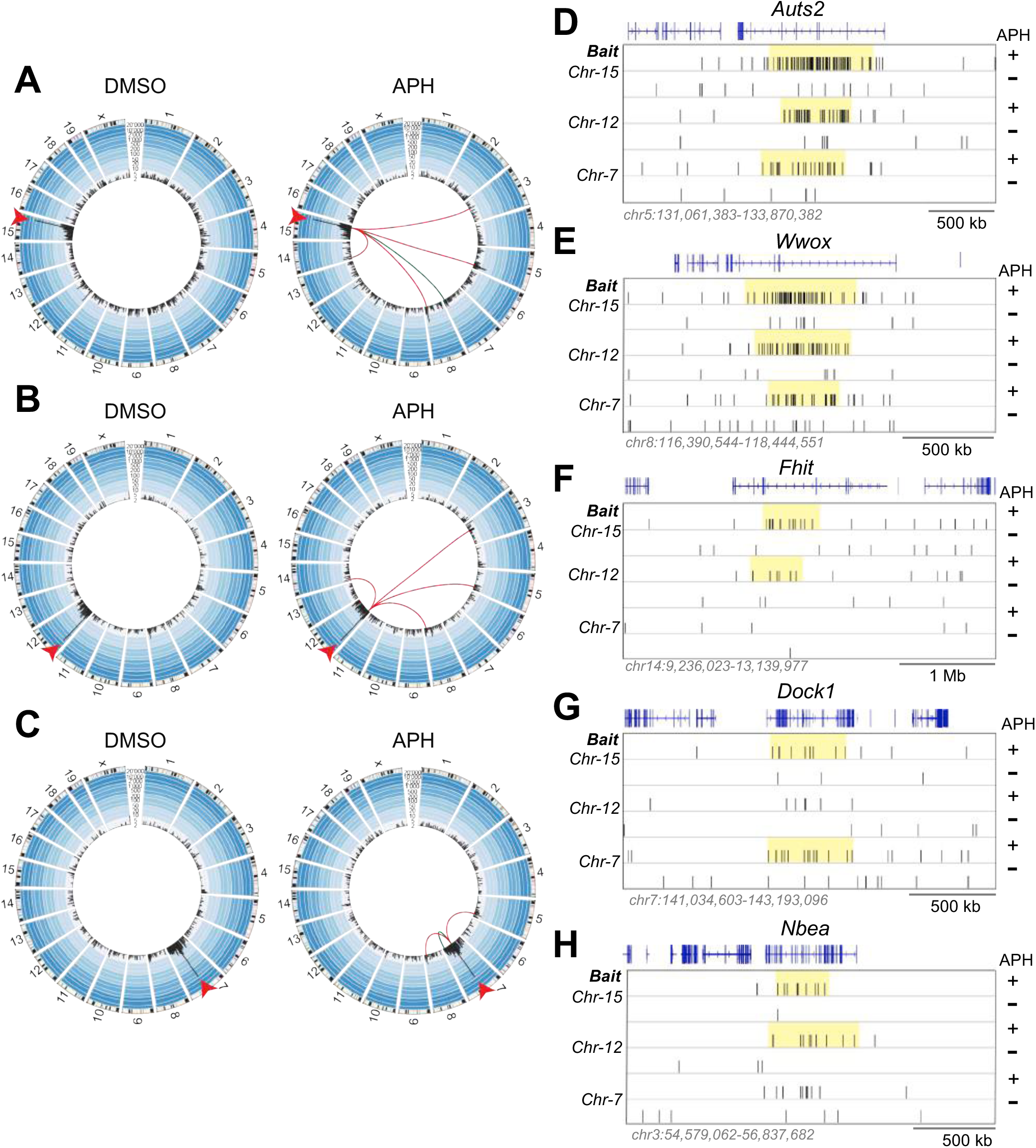
Genome-wide Identification of Replication Stress-induced RDC-genes in Embryonic Stem Cells Line 2. (***A-C***) Circos plots of the mouse genome divided into individual chromosomes showing the genome-wide LAM-HTGTS junction pattern of *Chr*-15, *Chr*-12, and *Chr*-7 sgRNA-mediated bait DSBs in the second *Xrcc4*^*−/−*^*p53*^*−/−*^ ESC line. Figures are organized as described in Figs. 1*A-C*. Red lines connect the break-site to 3 replication stress-induced RDCs identified by bait DSBs on all three tested chromosomes. Green lines connect the break-site to RDCs identified by two of three HTGTS baits. **(*D-H*)** 20,000 randomly selected LAM-HTGTS prey junctions from APH-(+) or DMSO-treated (-) experiments are plotted. Each panel represents the RDC-genes discovered by either two or three independent HTGTS-baits on the indicated chromosomes. The yellow rectangle indicates RDC location. RefGene (blue track) indicates the gene location.

**Figure S2.**
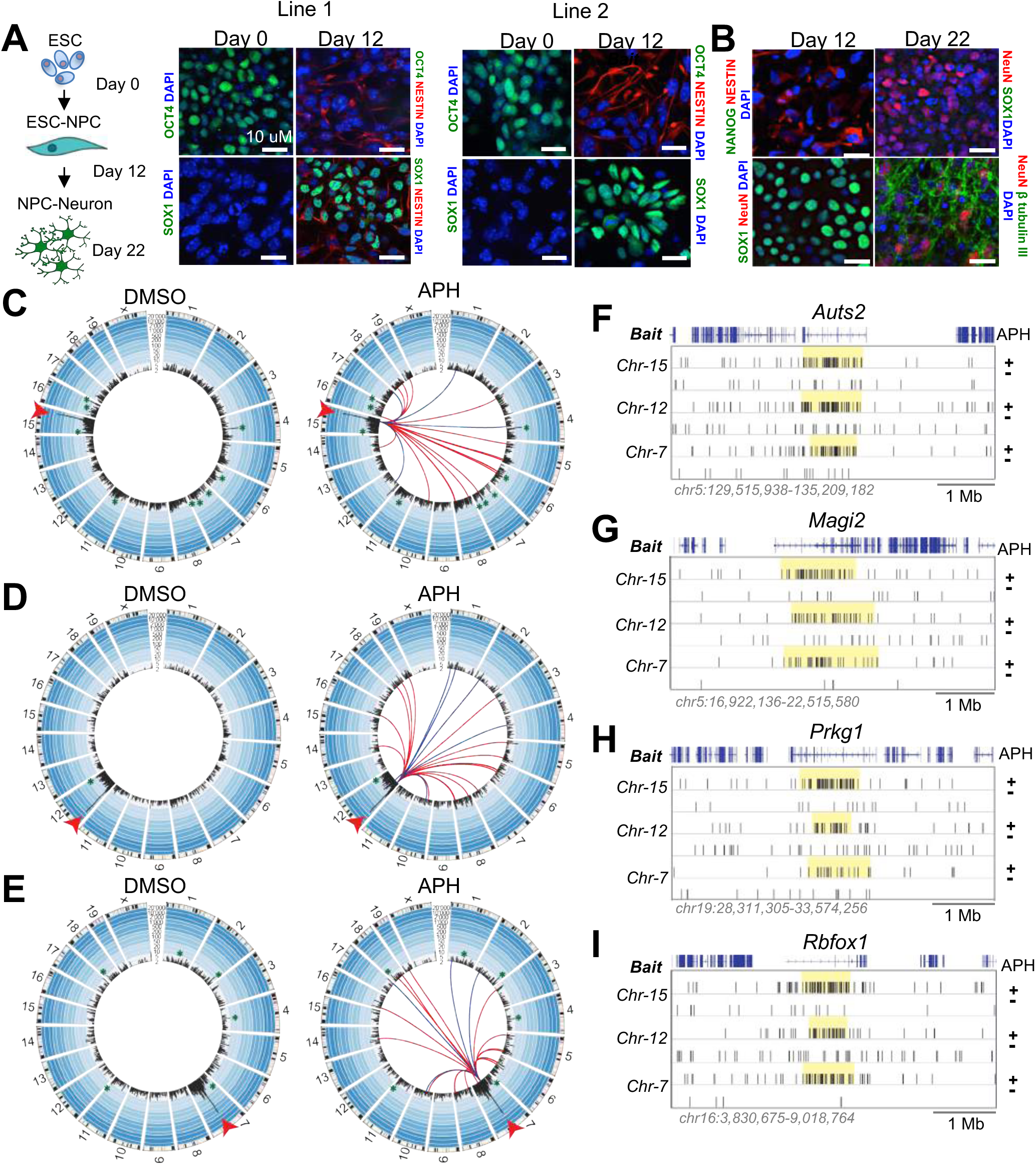
Robust RDC-genes Identified in a Second Line of Embryonic Stem Cells-Derived Neural Progenitor Cells. (***A***) *Left* panel: the experiment timeline of the *in vitro* ESC-derived NPC and neurons. *Middle* and *right* panels: immunofluorescence staining of *Xrcc4*^*−/−*^*p53*^*−/−*^ ESCs at day 0 and day 12 post NPC induction. Cells were stained for an ESC-specific marker *Oct4* and NPC-specific markers *Sox1* and *Pax6*. **(*B*)** Immunofluorescence staining of *Xrcc4*^*−/−*^*p53*^*−/−*^ neurons at indicated timepoint after differentiation. Cells were stained with NPC-specific markers *Nestin* and *Sox1*, an ESC-specific marker *Nanog*, and neuronal-specific markers *NeuN* and *β Tubulin III*. Nuclei were stained with DAPI (blue). **(*C-E*)** Circos plots showing genome-wide LAM-HTGTS junctions identified in the second line of *Xrcc4*^*−/−*^*p53*^*−/−*^ ESC-NPCs. Plots are organized as described in Figs. 2*A-C*. 10,000 randomly selected junctions from four independent experiments are plotted for DMSO- and APH-treated experiments. Red lines connect the break-site to 17 replication stress-induced RDCs identified by bait DSBs on all three tested chromosomes. Blue lines in each plot indicate RDCs identified by bait DSBs on two of the three tested break-sites, which numbered four for *Chr-15*, six for *Chr-12*, and four for *Chr-7* bait DSBs. **(*F-I*)** 20,000 randomly selected LAM-HTGTS prey junctions from APH-(+) or DMSO-treated (-) experiments are plotted for *Auts2, Magi2, Prkg1* and *Rbfox1* RDC-genes. Figures are organized as described in Fig. 2*D-H*.

**Figure S3.**
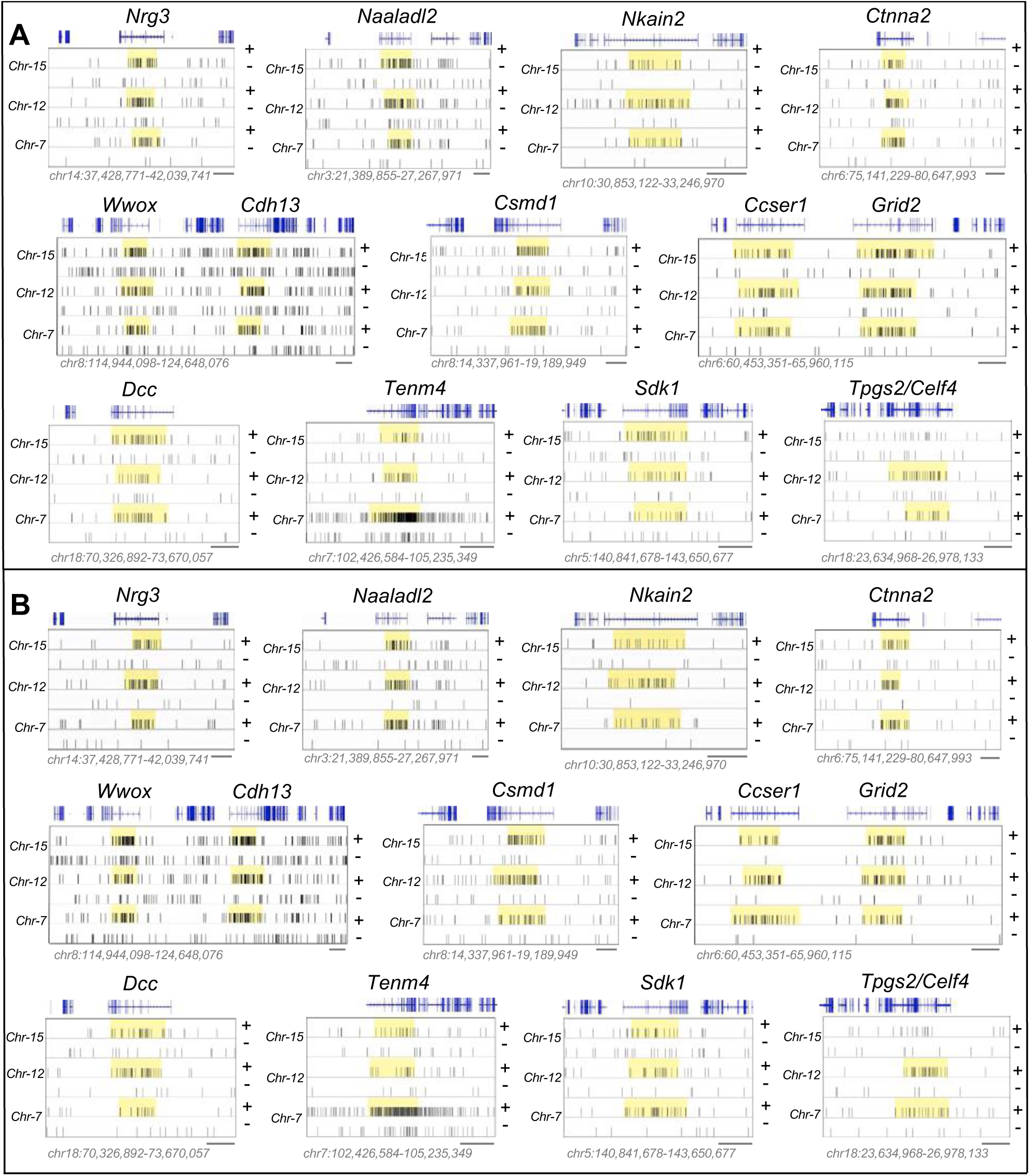
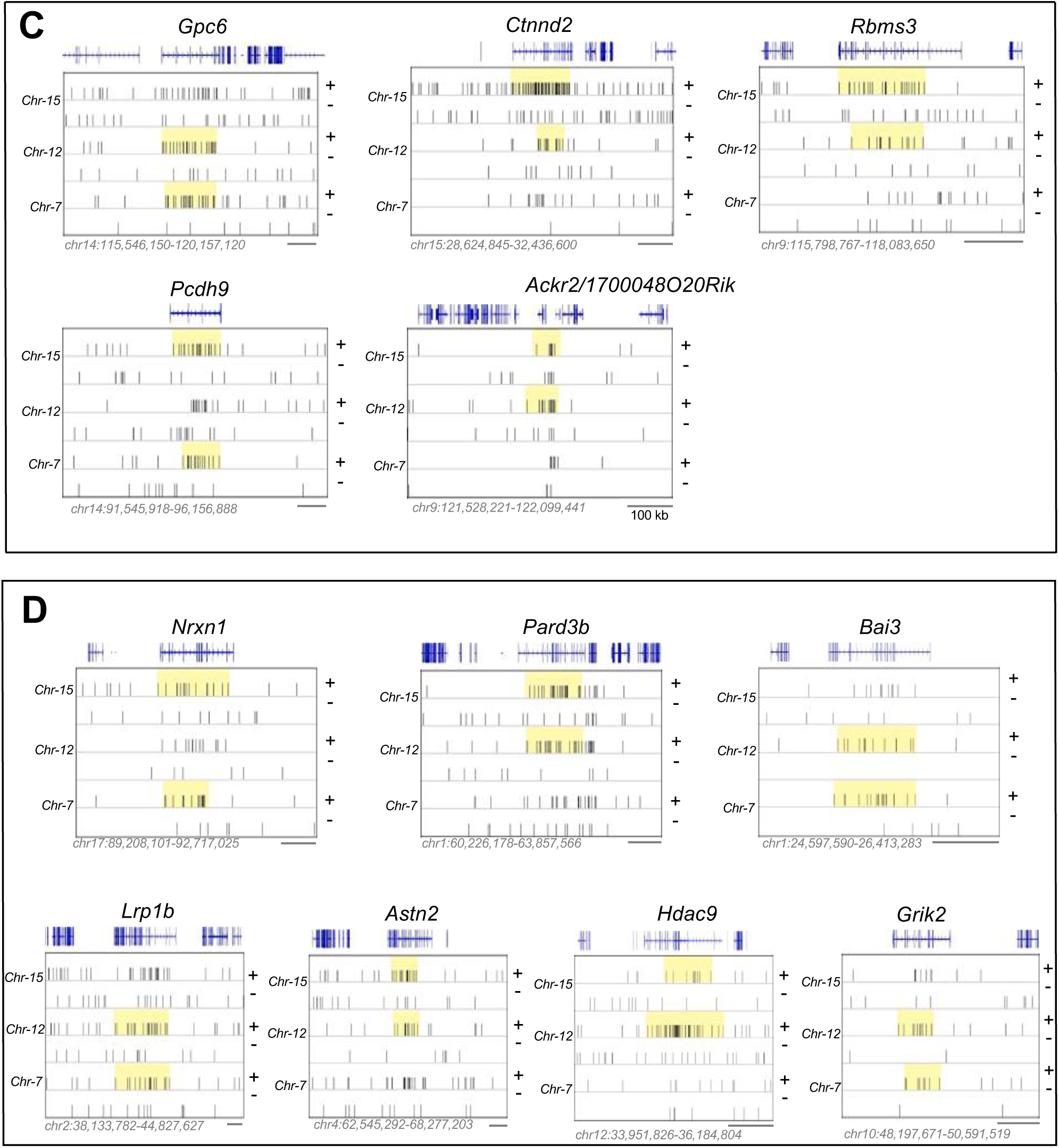
Shared and Unique RDC-genes in Differentiated Neural Progenitor Cells. LAM-HTGTS junction plots of shared RDC-genes in ESC-NPC line *1* (***A***) and line *2* (***B***), RDC-genes that are unique to line 1 (**C**), or RDC-genes that are unique to line 2 (**D**). Figures are organized as described in Figs. 1*D-H*. Scale bars: 500 kb, except 100 kb for the *Ackr2/1700048O20Rik* locus.

**Figure S4.**
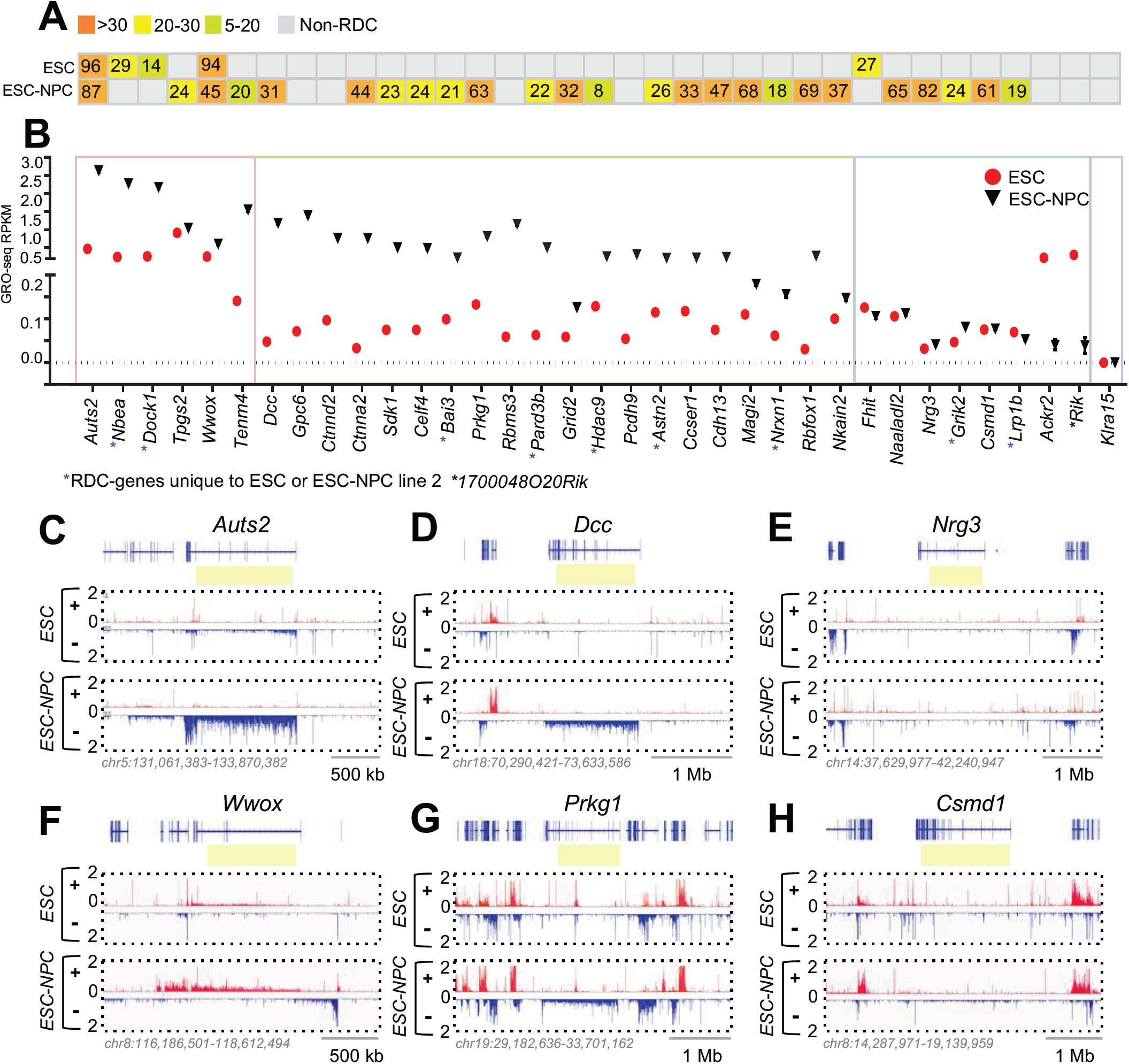
Transcription Activity in the Second Line of Embryonic Stem Cells and Neural Progenitor Cells. (***A***) RDC robustness score is indicated for each RDC-gene. (***B***) Transcription activity of RDC-genes. Figure is organized as described in Fig.3*B*. **(*C-H*)** Panels represent robust RDC-genes in ESCs and differentiated NPCs with different transcription levels in both cell lines. Figures are organized as described in Figure 3*C-H*.

## TABLES

**Table S1.**
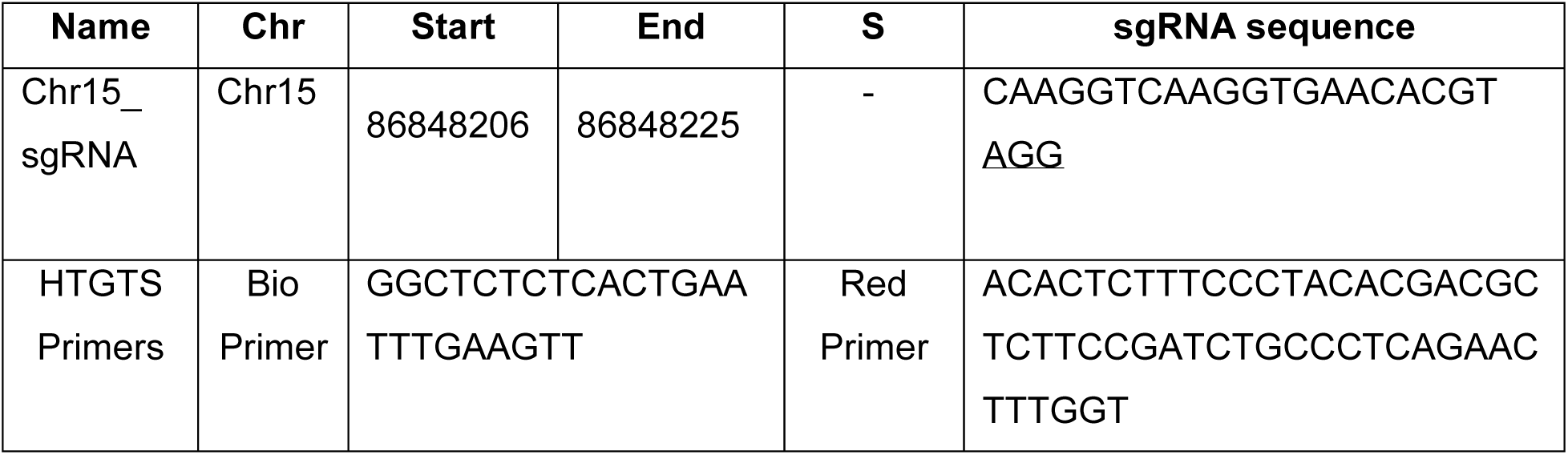
Chromosome-Specific, HTGTS Bait DSB-Inducing SgRNA, and HTGTS Primers Related to Figure 1. Sequence and coordinates of site-specific *Chr-15* LAM-HTGTS bait with the sgRNA. Biotinylated (Bio-primer) and the nested PCR (Red) primer sequences specific for CRISPR/Cas9-induced *Chr-15* HTGTS locus. For the sgRNA sequences and LAM-HTGTS primers used for *Chr-12*, and *-7*, see Wei et al., 2016; 2018. “Start” and “End” indicate the sgRNA targeting sequences. PAM motif is underlined. Chr: chromosome, S: strand.

**Table S2.**
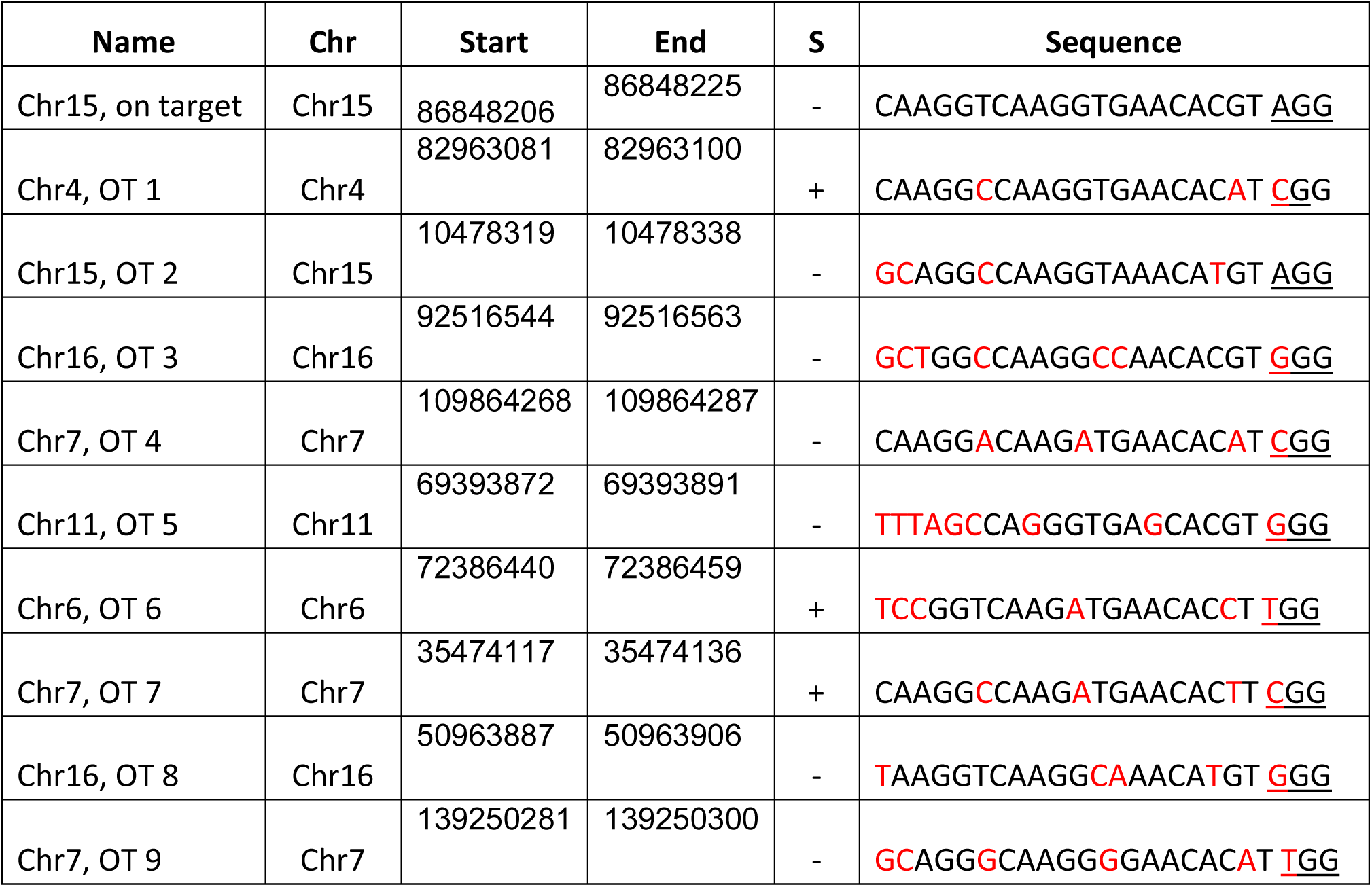
Off-Target Locations of HTGTS Bait-Inducing SgRNA for Chr-15, Related to Results. Table shows the sequences of the sgRNA on-target and off-target (OT) sites for Chr-15 sgRNA. For Chr-12-sgRNA-1 and Chr-7-sgRNA, please see Wei et al., 2016; 2018. Chr: chromosome; S: strand. Mismatch nucleotides between on-target and off-target sites are highlighted in red. PAM sequence is underlined.

**Table S-3.** LAM-HTGTS Unique Junction Numbers, Related to Figure 1 and 2 and Supplementary Figure 1 and 2. Table shows the number of unique LAM-HTGTS junctions of each experiment performed on *Xrcc4*^−/−^*p53*^−/−^ ESCs and *Xrcc4*^−/−^*p53*^−/−^ ESC-NPCs. Number of total junction (column 4), junctions given by joining the resected DSBs at ± 10 kb around breaksite (column 5), junctions ± 10 kb away from the bait DSBs at cis-chromosome (Intra-chromosomal; column 6), and junctions at non-breaksite containing chromosomal (Inter-chromosomal, column 7) from each experiment were shown.

| HTGTS Bait | Cell Type | Treatment | Total | Break site | Intra Chromosomal | Inter Chromosomal |
| --- | --- | --- | --- | --- | --- | --- |
| Chr-15-sgRNA | ESC-line 1 | DMSO-Exp A | 5225 | 3328 | 545 | 1352 |
| Chr-15-sgRNA | ESC-line 1 | DMSO-Exp B | 4733 | 3429 | 433 | 871 |
| Chr-15-sgRNA | ESC-line 1 | DMSO-Exp C | 5285 | 3269 | 510 | 1506 |
| Chr-15-sgRNA | ESC-line 1 | DMSO-Exp D | 6243 | 3783 | 586 | 1874 |
| Chr-15-sgRNA | ESC-line 1 | APH-Exp A | 8860 | 5577 | 990 | 2293 |
| Chr-15-sgRNA | ESC-line 1 | APH-Exp B | 8191 | 4902 | 904 | 2385 |
| Chr-15-sgRNA | ESC-line 1 | APH-Exp C | 5425 | 3462 | 501 | 1462 |
| Chr-15-sgRNA | ESC-line 1 | APH-Exp D | 8652 | 5539 | 960 | 2153 |
| Chr-12-sgRNA | ESC-line 1 | DMSO-Exp A | 3604 | 2774 | 310 | 520 |
| Chr-12-sgRNA | ESC-line 1 | DMSO-Exp B | 4337 | 3130 | 474 | 733 |
| Chr-12-sgRNA | ESC-line 1 | DMSO-Exp C | 5085 | 3598 | 525 | 962 |
| Chr-12-sgRNA | ESC-line 1 | DMSO-Exp D | 6097 | 3592 | 843 | 1662 |
| Chr-12-sgRNA | ESC-line 1 | APH-Exp A | 4150 | 3043 | 453 | 654 |
| Chr-12-sgRNA | ESC-line 1 | APH-Exp B | 8732 | 5465 | 1584 | 1683 |
| Chr-12-sgRNA | ESC-line 1 | APH-Exp C | 9457 | 5718 | 1692 | 2047 |
| Chr-12-sgRNA | ESC-line 1 | APH-Exp D | 6324 | 3158 | 1095 | 2071 |
| Chr-7-sgRNA | ESC-line 1 | DMSO-Exp A | 6797 | 4431 | 1071 | 1295 |
| Chr-7-sgRNA | ESC-line 1 | DMSO-Exp B | 3828 | 3000 | 435 | 393 |
| Chr-7-sgRNA | ESC-line 1 | DMSO-Exp C | 3983 | 3196 | 442 | 345 |
| Chr-7-sgRNA | ESC-line 1 | DMSO-Exp D | 3117 | 2448 | 332 | 337 |
| Chr-7-sgRNA | ESC-line 1 | DMSO-Exp E | 2257 | 1708 | 234 | 315 |
| Chr-7-sgRNA | ESC-line 1 | DMSO-Exp F | 2512 | 1748 | 279 | 485 |
| Chr-7-sgRNA | ESC-line 1 | DMSO-Exp G | 2083 | 1820 | 145 | 118 |
| Chr-7-sgRNA | ESC-line 1 | DMSO-Exp I | 2101 | 1821 | 152 | 128 |
| Chr-7-sgRNA | ESC-line 1 | APH-Exp A | 8502 | 5428 | 1731 | 1343 |
| Chr-7-sgRNA | ESC-line 1 | APH-Exp B | 4843 | 3349 | 605 | 889 |
| Chr-7-sgRNA | ESC-line 1 | APH-Exp C | 4413 | 3322 | 506 | 585 |
| Chr-7-sgRNA | ESC-line 1 | APH-Exp D | 3852 | 3055 | 430 | 367 |
| Chr-7-sgRNA | ESC-line 1 | APH-Exp E | 2045 | 1535 | 203 | 307 |
| Chr-7-sgRNA | ESC-line 1 | APH-Exp F | 3765 | 2975 | 428 | 362 |
| Chr-7-sgRNA | ESC-line 1 | APH-Exp G | 2209 | 1744 | 188 | 277 |
| Chr-7-sgRNA | ESC-line 1 | APH-Exp I | 1717 | 1413 | 134 | 170 |
| Chr-15-sgRNA | ESC-line 2 | DMSO-Exp A | 3159 | 2180 | 232 | 747 |
| Chr-15-sgRNA | ESC-line 2 | DMSO-Exp B | 5178 | 3347 | 493 | 1338 |
| Chr-15-sgRNA | ESC-line 2 | DMSO-Exp C | 3268 | 2283 | 255 | 730 |
| Chr-15-sgRNA | ESC-line 2 | DMSO-Exp D | 5256 | 3406 | 465 | 1385 |
| Chr-15-sgRNA | ESC-line 2 | APH-Exp A | 5783 | 3240 | 640 | 1903 |
| Chr-15-sgRNA | ESC-line 2 | APH-Exp B | 6308 | 3872 | 648 | 1788 |
| Chr-15-sgRNA | ESC-line 2 | APH-Exp C | 4615 | 2948 | 390 | 1277 |
| Chr-15-sgRNA | ESC-line 2 | APH-Exp D | 5448 | 3510 | 538 | 1400 |
| Chr-12-sgRNA | ESC-line 2 | DMSO-Exp A | 4993 | 3266 | 799 | 928 |
| Chr-12-sgRNA | ESC-line 2 | DMSO-Exp B | 5351 | 3550 | 939 | 862 |
| Chr-12-sgRNA | ESC-line 2 | DMSO-Exp C | 4993 | 3310 | 772 | 911 |
| Chr-12-sgRNA | ESC-line 2 | DMSO-Exp D | 5348 | 3599 | 934 | 815 |
| Chr-12-sgRNA | ESC-line 2 | APH-Exp A | 9391 | 5154 | 2067 | 2170 |
| Chr-12-sgRNA | ESC-line 2 | APH-Exp B | 5111 | 3434 | 846 | 831 |
| Chr-12-sgRNA | ESC-line 2 | APH-Exp C | 5190 | 3332 | 893 | 965 |
| Chr-12-sgRNA | ESC-line 2 | APH-Exp D | 9197 | 5389 | 1949 | 1859 |
| Chr-7-sgRNA | ESC-line 2 | DMSO-Exp A | 5506 | 3449 | 847 | 1210 |
| Chr-7-sgRNA | ESC-line 2 | DMSO-Exp B | 3914 | 2679 | 554 | 681 |
| Chr-7-sgRNA | ESC-line 2 | DMSO-Exp C | 5980 | 3849 | 1045 | 1086 |
| Chr-7-sgRNA | ESC-line 2 | DMSO-Exp D | 7989 | 4995 | 1589 | 1405 |
| Chr-7-sgRNA | ESC-line 2 | APH-Exp A | 6262 | 4123 | 1190 | 949 |
| Chr-7-sgRNA | ESC-line 2 | APH-Exp B | 8314 | 5165 | 1699 | 1450 |
| Chr-7-sgRNA | ESC-line 2 | APH-Exp C | 5968 | 3866 | 1074 | 1028 |
| Chr-7-sgRNA | ESC-line 2 | APH-Exp D | 8477 | 5289 | 1723 | 1465 |
| Chr-15-sgRNA | ESC-NPC line 1 | DMSO-Exp A | 6689 | 3011 | 715 | 2963 |
| Chr-15-sgRNA | ESC-NPC line 1 | DMSO-Exp B | 7368 | 3421 | 1148 | 2799 |
| Chr-15-sgRNA | ESC-NPC line 1 | DMSO-Exp C | 6646 | 3125 | 643 | 2878 |
| Chr-15-sgRNA | ESC-NPC line 1 | DMSO-Exp D | 7560 | 3008 | 1352 | 3200 |
| Chr-15-sgRNA | ESC-NPC line 1 | APH-Exp A | 6634 | 3009 | 658 | 2967 |
| Chr-15-sgRNA | ESC-NPC line 1 | APH-Exp B | 7525 | 2968 | 1295 | 3262 |
| Chr-15-sgRNA | ESC-NPC line 1 | APH-Exp C | 6813 | 3046 | 681 | 3086 |
| Chr-15-sgRNA | ESC-NPC line 1 | APH-Exp D | 7643 | 2979 | 1380 | 3284 |
| Chr-12-sgRNA | ESC-NPC line 1 | DMSO-Exp A | 6226 | 3835 | 616 | 1775 |
| Chr-12-sgRNA | ESC-NPC line 1 | DMSO-Exp B | 8060 | 4135 | 1434 | 2491 |
| Chr-12-sgRNA | ESC-NPC line 1 | DMSO-Exp C | 6820 | 3188 | 991 | 2641 |
| Chr-12-sgRNA | ESC-NPC line 1 | DMSO-Exp D | 7271 | 4096 | 1150 | 2025 |
| Chr-12-sgRNA | ESC-NPC line 1 | APH-Exp A | 6842 | 3974 | 1126 | 1742 |
| Chr-12-sgRNA | ESC-NPC line 1 | APH-Exp B | 5427 | 3781 | 492 | 1154 |
| Chr-12-sgRNA | ESC-NPC line 1 | APH-Exp C | 8344 | 3447 | 1571 | 3326 |
| Chr-12-sgRNA | ESC-NPC line 1 | APH-Exp D | 6770 | 3662 | 677 | 2431 |
| Chr-7-sgRNA | ESC-NPC line 1 | DMSO-Exp A | 5979 | 3896 | 654 | 1429 |
| Chr-7-sgRNA | ESC-NPC line 1 | DMSO-Exp B | 7370 | 3413 | 1901 | 2056 |
| Chr-7-sgRNA | ESC-NPC line 1 | DMSO-Exp C | 6085 | 3857 | 690 | 1538 |
| Chr-7-sgRNA | ESC-NPC line 1 | DMSO-Exp D | 7366 | 3496 | 1798 | 2072 |
| Chr-7-sgRNA | ESC-NPC line 1 | APH-Exp A | 6323 | 3904 | 863 | 1556 |
| Chr-7-sgRNA | ESC-NPC line 1 | APH-Exp B | 7024 | 3838 | 1314 | 1872 |
| Chr-7-sgRNA | ESC-NPC line 1 | APH-Exp C | 5574 | 3735 | 564 | 1275 |
| Chr-7-sgRNA | ESC-NPC line 1 | APH-Exp D | 7534 | 3571 | 1782 | 2181 |
| Chr-15-sgRNA | ESC-NPC line 2 | DMSO-Exp A | 6534 | 2983 | 713 | 2838 |
| Chr-15-sgRNA | ESC-NPC line 2 | DMSO-Exp B | 7314 | 3430 | 1200 | 2684 |
| Chr-15-sgRNA | ESC-NPC line 2 | DMSO-Exp C | 6611 | 3025 | 664 | 2920 |
| Chr-15-sgRNA | ESC-NPC line 2 | DMSO-Exp D | 7538 | 2947 | 1361 | 3230 |
| Chr-15-sgRNA | ESC-NPC line 2 | APH-Exp A | 6694 | 3061 | 665 | 2968 |
| Chr-15-sgRNA | ESC-NPC line 2 | APH-Exp B | 6837 | 3025 | 694 | 3118 |
| Chr-15-sgRNA | ESC-NPC line 2 | APH-Exp C | 7620 | 4277 | 1330 | 3343 |
| Chr-15-sgRNA | ESC-NPC line 2 | APH-Exp D | 8250 | 3123 | 1327 | 3800 |
| Chr-12-sgRNA | ESC-NPC line 2 | DMSO-Exp A | 5434 | 3598 | 492 | 1344 |
| Chr-12-sgRNA | ESC-NPC line 2 | DMSO-Exp B | 7658 | 3507 | 1654 | 2497 |
| Chr-12-sgRNA | ESC-NPC line 2 | DMSO-Exp C | 5655 | 3607 | 556 | 1492 |
| Chr-12-sgRNA | ESC-NPC line 2 | DMSO-Exp D | 7668 | 3483 | 1654 | 2504 |
| Chr-12-sgRNA | ESC-NPC line 2 | APH-Exp A | 5104 | 3371 | 488 | 1245 |
| Chr-12-sgRNA | ESC-NPC line 2 | APH-Exp B | 6872 | 3568 | 1258 | 2046 |
| Chr-12-sgRNA | ESC-NPC line 2 | APH-Exp C | 6737 | 3928 | 807 | 2002 |
| Chr-12-sgRNA | ESC-NPC line 2 | APH-Exp D | 7056 | 3964 | 1229 | 1863 |
| Chr-7-sgRNA | ESC-NPC line 2 | DMSO-Exp A | 6009 | 3633 | 832 | 1544 |
| Chr-7-sgRNA | ESC-NPC line 2 | DMSO-Exp B | 7160 | 2902 | 2161 | 2097 |
| Chr-7-sgRNA | ESC-NPC line 2 | DMSO-Exp C | 5993 | 3552 | 820 | 1621 |
| Chr-7-sgRNA | ESC-NPC line 2 | DMSO-Exp D | 7746 | 2914 | 2353 | 2479 |
| Chr-7-sgRNA | ESC-NPC line 2 | APH-Exp A | 6134 | 3685 | 753 | 1696 |
| Chr-7-sgRNA | ESC-NPC line 2 | APH-Exp B | 7197 | 3005 | 2213 | 1979 |
| Chr-7-sgRNA | ESC-NPC line 2 | APH-Exp C | 5724 | 3523 | 751 | 1450 |
| Chr-7-sgRNA | ESC-NPC line 2 | APH-Exp D | 6809 | 2748 | 1981 | 2080 |

**Table S4:** Complete List of RDC-genes Discovered in Embryonic Stem Cells and Differentiated Neural Progenitor Cells. All RDCs discovered in two ESC lines and their differentiated NPCs are sorted based on gene length. Average RPKM values derived from three independent GRO-seq experiments are shown (columns 4-7). Orange boxes highlight the RDC-genes that were discovered in the corresponding cell types. Genes marked with an asterisk, (*Tpgs2/Celf4* and *Ackr2/1700048O20Rik*) are nearby small genes encompassing an RDC area. All other RDCs were enriched in single genes. Black asterisk: RDC in both ESC-NPC lines. Red asterisk: RDC only in ESC-NPC line 1. Replication timing score (R.T.) of all RDC-genes is shown at the last column. Chr: chromosome.

| Chr | Start | End | Gene | Length (Mb) | ESC_1 RPKM | ESC_2 RPKM | NPC_1 RPKM | NPC_2 RPKM | R.T. |
| --- | --- | --- | --- | --- | --- | --- | --- | --- | --- |
| chr2 | 40452292 | 42509068 | *Lrp1b* | 2.056 | 0.05 | 0.07 | 0.11 | 0.05 | Late |
| chr8 | 15892544 | 17535335 | *Csmd1* | 1.642 | 0.04 | 0.08 | 0.15 | 0.08 | Late |
| chr14 | 10382607 | 11994496 | *Fhit* | 1.611 | 0.11 | 0.13 | 0.07 | 0.11 | Late |
| chr16 | 5884935 | 7412573 | *Rbfox1* | 1.527 | 0.04 | 0.03 | 0.23 | 0.28 | Late |
| chr5 | 18732913 | 20210609 | *Magi2* | 1.477 | 0.10 | 0.11 | 0.23 | 0.18 | Late |
| chr6 | 63206900 | 64616273 | *Grid2* | 1.409 | 0.04 | 0.05 | 0.27 | 0.13 | Late |
| chr3 | 23798103 | 25144144 | *Naaladl*<br>*2* | 1.346 | 0.09 | 0.11 | 0.10 | 0.11 | Late |
| chr6 | 61130368 | 62332857 | *Ccser1* | 1.202 | 0.07 | 0.12 | 0.27 | 0.22 | Late |
| chr10 | 31409124 | 32609671 | *Nkain2* | 1.200 | 0.08 | 0.10 | 0.16 | 0.15 | Late |
| chr19 | 30638976 | 31839473 | *Prkg1* | 1.200 | 0.16 | 0.13 | 0.29 | 0.81 | Late |
| chr5 | 131913551 | 133018163 | *Auts2* | 1.104 | 0.64 | 0.47 | 2.25 | 2.63 | Neutral |
| chr14 | 39182136 | 40286326 | *Nrg3* | 1.104 | 0.03 | 0.03 | 0.14 | 0.04 | Late |
| chr6 | 76831630 | 77929611 | *Ctnna2* | 1.098 | 0.05 | 0.03 | 0.88 | 0.76 | Late |
| chr18 | 71413285 | 72510673 | *Dcc* | 1.097 | 0.03 | 0.05 | 2.02 | 1.19 | Late |
| chr17 | 90432983 | 91492092 | *Nrxn1* | 1.059 | 0.06 | 0.06 | 0.39 | 0.16 | Late |
| chr9 | 116481863 | 117538981 | *Rbms3* | 1.057 | 0.18 | 0.06 | 0.54 | 1.16 | Late |
| chr14 | 117324568 | 118378751 | *Gpc6* | 1.054 | 0.18 | 0.07 | 1.16 | 1.39 | Early/Late |
| chr8 | 120807704 | 121847348 | *Cdh13* | 1.039 | 0.14 | 0.07 | 0.22 | 0.24 | Early/Late |
| chr4 | 65041836 | 66065467 | *Astn2* | 1.023 | 0.09 | 0.11 | 0.35 | 0.22 | Late |
| chr1 | 61685447 | 62688858 | *Pard3b* | 1.003 | 0.05 | 0.06 | 0.78 | 0.50 | Early/Late |
| chr5 | 141717537 | 142689745 | *Sdk1* | 0.972 | 0.08 | 0.07 | 1.00 | 0.52 | Late |
| chr8 | 116963601 | 117876612 | *Wwox* | 0.913 | 0.43 | 0.26 | 0.37 | 0.63 | Neutral |
| chr14 | 93412918 | 94289838 | *Pcdh9* | 0.876 | 0.05 | 0.05 | 0.28 | 0.33 | Late |
| chr12 | 34732446 | 35601717 | *Hdac9* | 0.869 | 0.09 | 0.13 | 0.66 | 0.27 | Late |
| chr15 | 30102397 | 30959098 | *Ctnnd2* | 0.856 | 0.08 | 0.10 | 1.31 | 1.27 | Late |
| chr1 | 25124320 | 25886502 | *Bai3* | 0.762 | 0.12 | 0.09 | 2.19 | 2.24 | Late |
| chr7 | 103359196 | 104057064 | *Tenm4* | 0.697 | 0.63 | 0.14 | 1.60 | 1.60 | Early |
| chr10 | 48819268 | 49508510 | *Grik2* | 0.689 | 0.03 | 0.05 | 0.11 | 0.12 | Early |
| chr3 | 55429119 | 55987573 | *Nbea* | 0.558 | 0.34 | 0.25 | 2.54 | 2.30 | Early |
| chr7 | 141862419 | 142365330 | *Dock1* | 0.502 | 0.90 | 0.27 | 2.21 | 2.18 | Early |
| chr18 | 25636120 | 25912434 | *Celf4\** | 0.276 | 0.07 | 0.07 | 1.08 | 1.05 | Early |
| chr18 | 25285723 | 25327328 | *Tpgs2\** | 0.041 | 0.59 | 0.86 | 1.67 | 1.63 | Early |
| chr9 | 121807522 | 121820189 | *Ackr2\** | 0.012 | 0.05 | 0.25 | 0.03 | 0.05 | Early |
| chr9 | 121846442 | 121856134 | *170004*<br>*8O20Ri*<br>*k\** | 0.009 | 0.09 | 0.31 | 0.13 | 0.02 | Early |

**Dataset S1, Supplemented as Excel file:**

The dataset provides information about RDCs identified in ESCs and ESC-NPCs. The SICER-determined RDC coordinates (columns A to C), the group of given RDC (column D), the genes that are greater than 80 kb within given RDCs (column E), the location of chromosomal bait DSBs that identified given RDCs (column F), the RDC robustness score (column G), and the false discovery rate (FDR) adjusted, MACS-based P-value of each RDC (columns H to K) for line 1 and line 2 are provided. The dataset is ranked by the RDC robustness score (column G). The robust RDCs (RSS > 30) identified in both lines at either ESC or ESC-NPC status are highlighted in green. Chrom: chromosome. Cis-bait: P-values of RDCs identified by the HTGTS bait at the *cis*-chromosome. Trans-bait: P-values of RDCs identified by the HTGTS bait at the *trans*-chromosomes.

**Dataset S2, Supplemented as Excel file:**

The dataset provides information about RDC candidates identified by the *Chr-7, Chr-12*, or *Chr-15* HTGTS-baits in both ESC lines. The coordinates of RDC candidate (column A-C), the FDR adjusted MACS-based P-value (column D), and genes that are greater than 80 kb in given RDCs (column E) are shown. We highlighted RDC candidates that were identified as RDCs in the other line (green or blue) or in the primary NSPCs (orange). Chrom: chromosome. Start/End: SICER-determined RDC-candidate location. Gene: Name of the RDC-gene or group of genes where HTGTS junctions were enriched.

**Dataset S3, Supplemented as Excel file:**

The dataset shows the information about RDCs candidates identified by the *Chr-7, Chr-12*, or *Chr-15* HTGTS-baits in two ESC-NPC lines. Table is organized as described in Dataset S2. Some RDCs that were identified in ESC-NPC line 1 were RDC-candidates in ESC-NPC line 2. They are highlighted in green in the detected bait chromosomes. Furthermore, some RDCs identified in ESC-NPC line 2 were RDC-candidates in ESC-NPC line 1. They are also highlighted in green in the bait-chromosomes that detected them. RDCs or RDC-candidates that were identified in primary NSPCs are RDC-candidates in either ESC-NPC line 1 or line 2. They are highlighted in orange. Yellow color highlights an ESC line 2 RDC that is an RDC-candidate in both ESC-NPC lines.

**Dataset 1.**
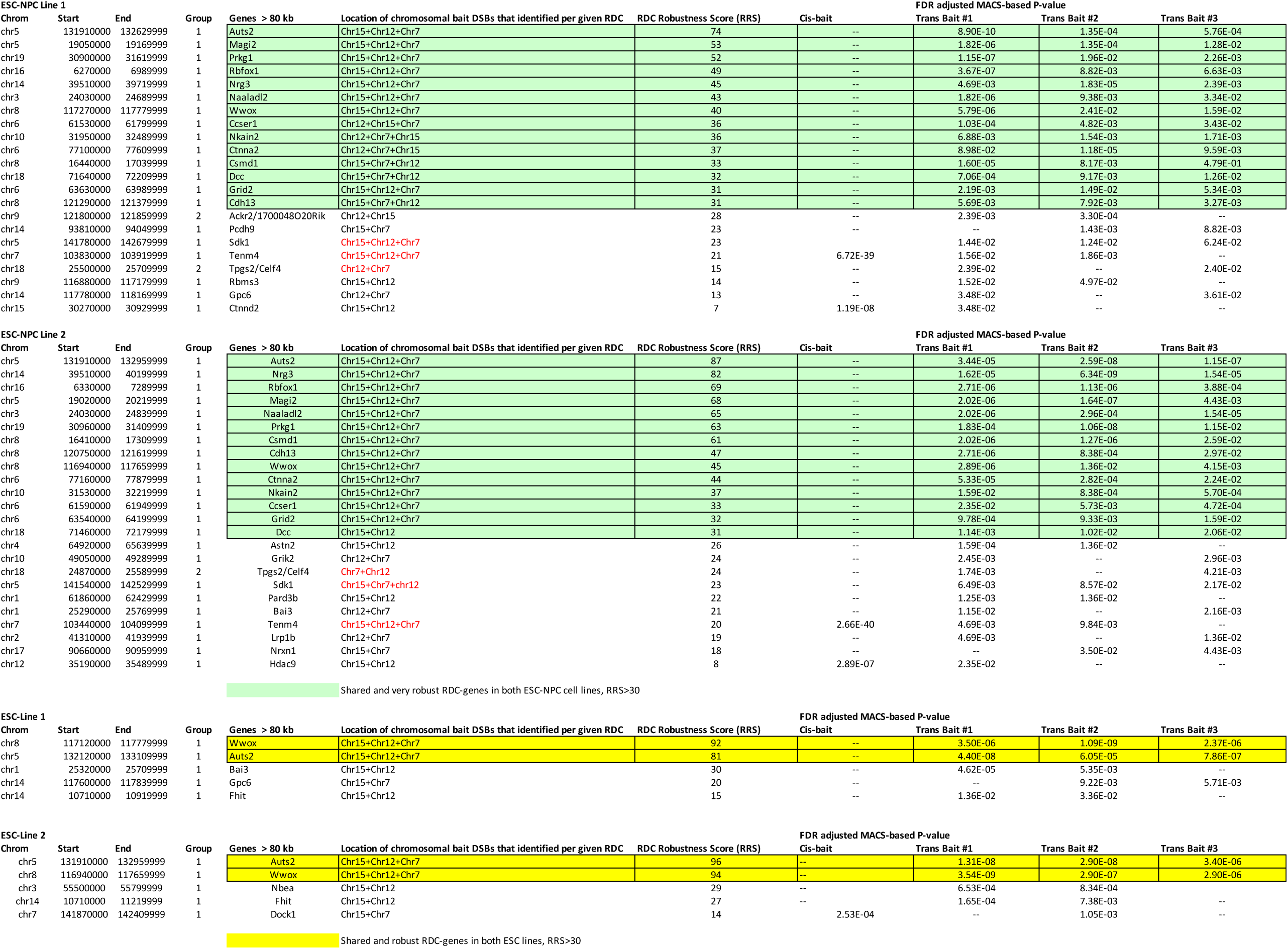

**Dataset 2.**
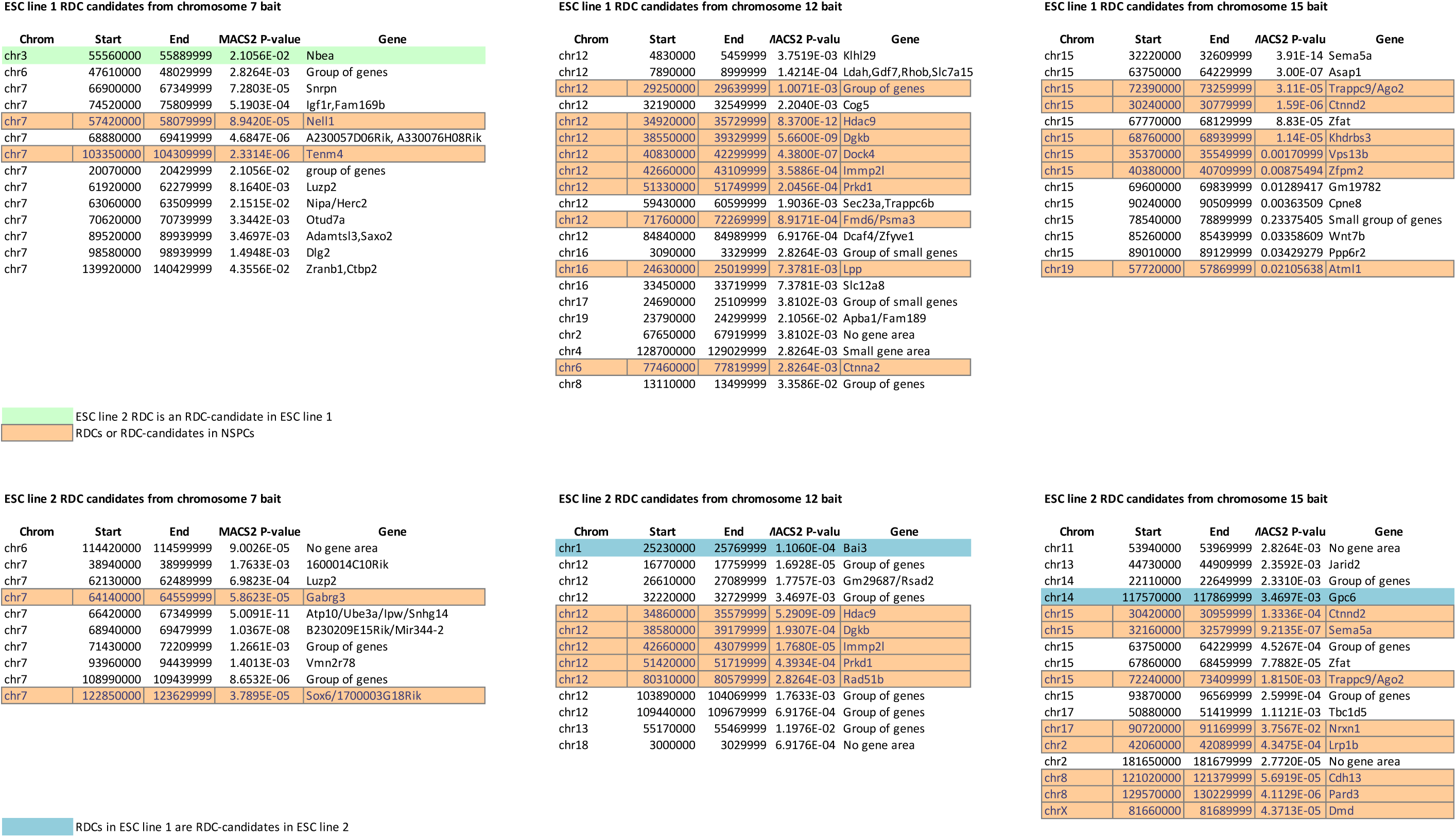

**Dataset 3.**
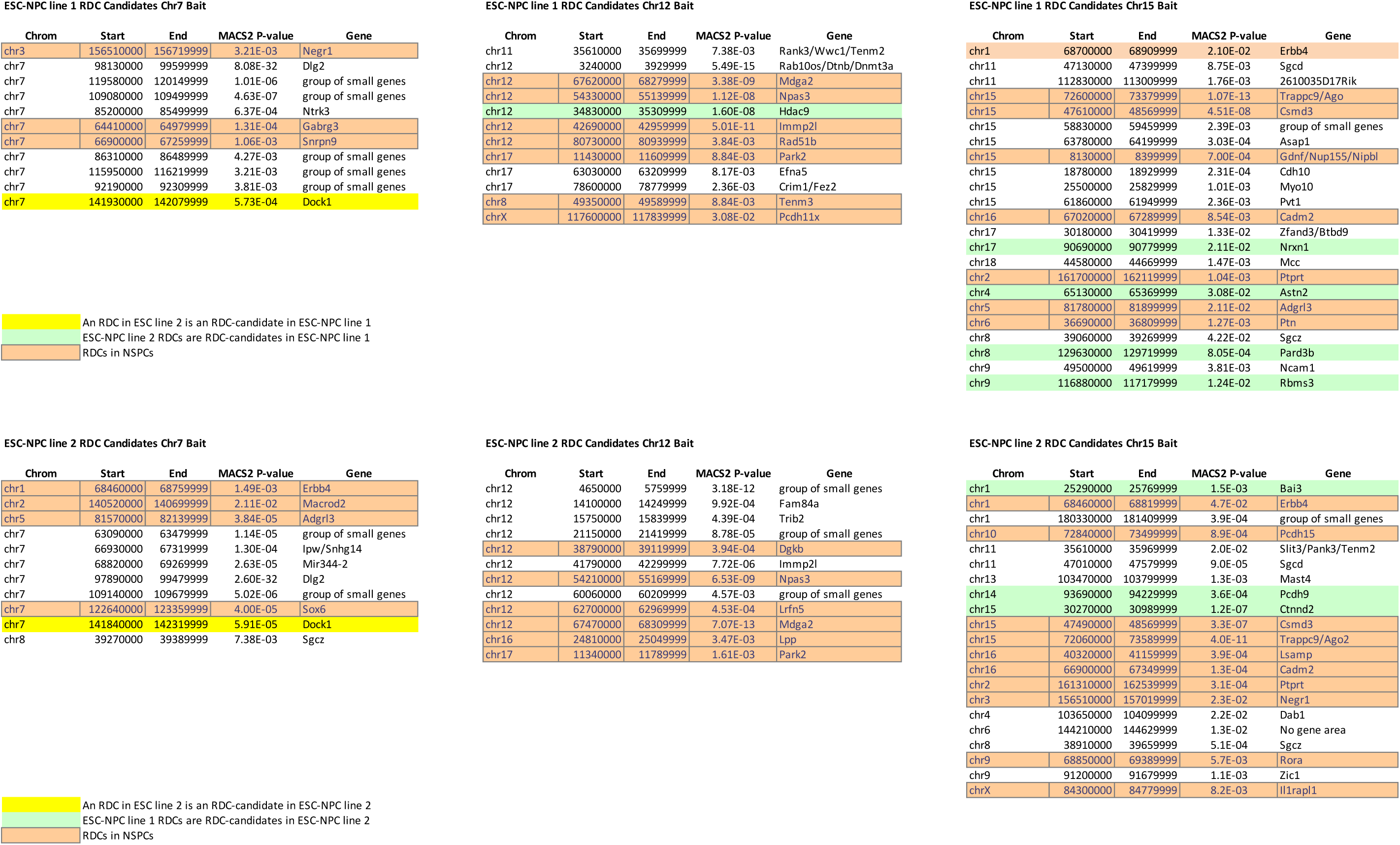

